# Multi-view confocal microscopy enables multiple organ and whole organism live-imaging

**DOI:** 10.1101/2021.05.04.442565

**Authors:** Olivier Leroy, Eric van Leen, Philippe Girard, Aurélien Villedieu, Christian Hubert, Floris Bosveld, Yohanns Bellaïche, Olivier Renaud

## Abstract

Understanding how development is coordinated in multiple tissues and gives rise to fully functional organs or whole organisms necessitates microscopy tools. Over the last decade numerous advances have been made in live-imaging, enabling high resolution imaging of whole organisms at cellular resolution. Yet, these advances mainly rely on mounting the specimen in agarose or aqueous solutions, precluding imaging of organisms whose oxygen uptake depends on ventilation. Here, we implemented a multi-view multi-scale microscopy strategy based on confocal spinning disk microscopy, called Multi-View confocal microScopy (MuViScopy). MuViScopy enables live-imaging of multiple organs with cellular resolution using sample rotation and confocal imaging without the need of sample embedding. We illustrate the capacity of MuViScopy by live-imaging *Drosophila melanogaster* pupal development throughout metamorphosis, highlighting how internal organs are formed and multiple organ development is coordinated. We foresee that MuViScopy will open the path to better understand developmental processes at the whole organism scale in living systems that necessitates gas exchange by ventilation.

## Introduction

The size, shape and organization of organs, tissues or cells are highly regulated during development, homeostasis and repair. As such, capturing cell, tissue and organ growth and morphogenesis over long periods of time, is central to understand these biological phenomena and to probe their underlying genetic and biophysical regulations (Collinet and Lecuit, 2021; Goodwin and Nelson, 2021; Hannezo and Heisenberg, 2019). Importantly, cell, tissue or organ dynamics can be driven by long range mechanical coupling, as well as systemic hormonal regulation (Barresi and Gilbert, 2019; Boulan et al., 2015; Cole et al., 2019; Villedieu et al., 2020), highlighting that the understanding of growth and morphogenesis also necessitates cellular imaging of multiple tissues or organs in whole living organisms.

Fluorescent light-microscopy has emerged as an essential technology to study biological systems by live imaging at spatiotemporal resolutions ranging from subcellular to whole organism scale (Keller, 2013). This is best illustrated by two distinct imaging approaches: confocal microscopy and light-sheet microscopy (Fig. 1A) (Bayguinov et al., 2018; Keller and Stelzer, 2008; Keller et al., 2008; Oreopoulos et al., 2014; Wan et al., 2019). Confocal fluorescence microscopes are currently the most commonly used imaging systems with optical sectioning capability. In particular, spinning disk confocal microscopy is a mature, commercially available technology for studying development, homeostasis and repair by live imaging. It can resolve biological processes from the subcellular to whole-tissue scales. Furthermore, spinning disk confocal microscopy has many advantages over laser scanning confocal microscopy for dynamic imaging, particularly in terms of acquisition speed, photobleaching and phototoxicity (Oreopoulos et al., 2014; Wang et al., 2005). Spinning disk confocal microscopy samples are often mounted on glass slides or specific dishes, where observation is only possible from one side, namely perpendicular to the slide. For many applications in biology, and in particular in the field of developmental biology, it is critical to be able to obtain multiple views of the structure(s) under investigation. Indeed, the observed tissues are often not organized in a uniform layer but present complex 3D structures. In confocal microscopy at the level of individual cells, attempts have been made to better orientate cells by inserting single cells into a glass capillary or fixing them to a glass fiber (Bradl et al., 1994; Bruns et al., 2015; Staier et al., 2011). Whether these approaches would allow to perform a multi-view exploration of tissues or organisms while achieving high resolution imaging of multiple tissues remains unexplored. Complementing confocal microscopy, light-sheet fluorescence microscopy (LSFM) combines intrinsic optical sectioning with wide-field detection (Keller, 2013). In contrast to epifluorescence microscopy only a thin slice (usually a few hundred nanometers to a few micrometers) of the sample is illuminated perpendicularly to the direction of observation. This method allows the acquisition of large field of view images in a single, spatially confined illumination step; offering low photobleaching, low phototoxicity, good signal-to-noise ratio and improved acquisition speed (de Medeiros et al., 2015; Guignard et al., 2020; Huisken, 2004; McDole et al., 2018; Mertz, 2011; Pitrone et al., 2013; Siedentopf and Zsigmondy, 1902; Strnad et al., 2016; Voie et al., 1993). In addition, the majority of light-sheet microscopes enable sample imaging from different points of view using a sample holder mounted on a rotating stage and/or by having multiple lenses to collect light from different angles (Chhetri et al., 2015; Krzic et al., 2012; Schmid et al., 2013; Tomer et al., 2012; Wu et al., 2013). So far, LSFM permits live *in toto* imaging of immersed/embedded samples. However, there are numerous model organisms that cannot be immersed or embedded as their survival depends on oxygen uptake by ventilation. Whether LSFM can be efficiently applied to non-immersed samples has yet to be tested.

**Figure 1.**
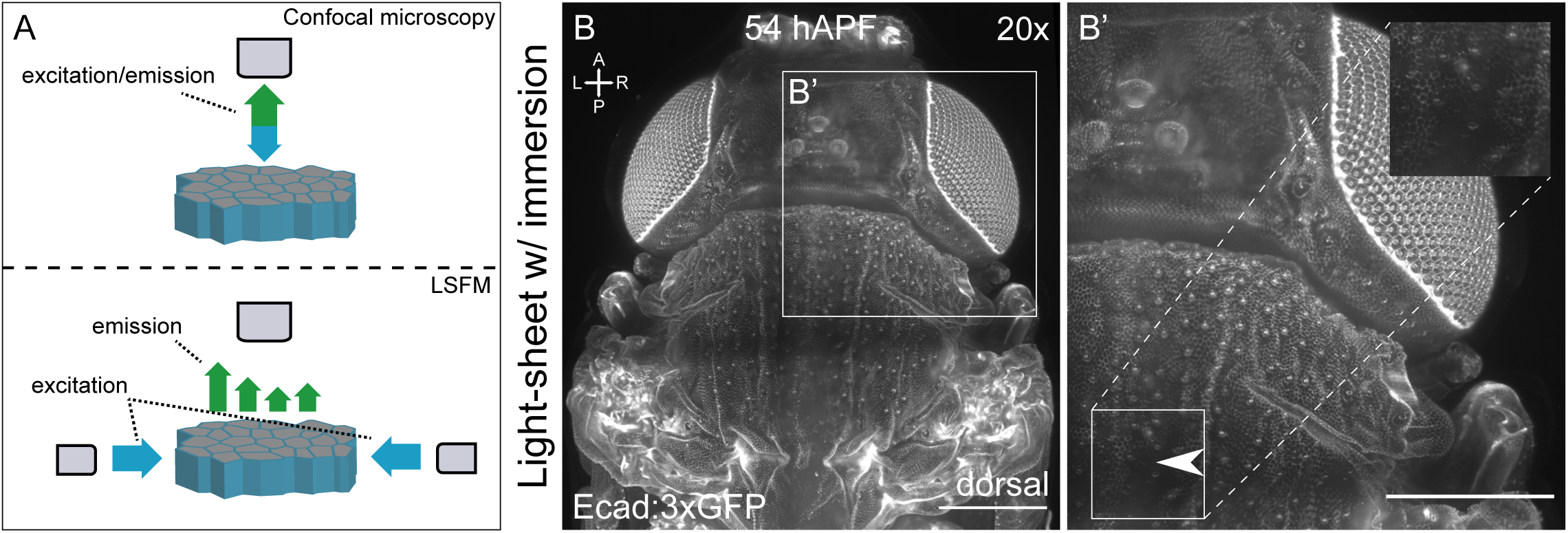
Light-sheet microscopy imaging of an immersed *Drosophila* pupa. A) Schematic of the difference in illumination/emission paths between dual-side illumination LSFM and confocal microscopy. In LSFM the illumination objective and the capturing objective are positioned at a 90° angle, while in confocal microscopy the same objective is used. B) Dorsal view of Ecad:3xGFP pupa imaged immersed in PBS using the Zeiss Z1 light-sheet microscope at 20x using dual-side illumination. The Anterior-Posterior (A-P) and Left-Right (L-R) axes are indicated. B’) Close-up of the region outlined in (B) as well as close-up in the white square region (insert) where the image quality is lowest (arrowhead).

As exemplified by numerous past and recent studies, the imaging *Drosophila* pupa without immersion has been key in the exploration of developmental and repair processes and to characterize core and conserved genetic and biophysical mechanisms driving cell differentiation as well as tissue proliferation, shaping, architecture and repair (Aigouy et al., 2010; Arata et al., 2017; Corson et al., 2017; Curran et al., 2017; Diaz-de-la-Loza et al., 2018; Etournay et al., 2015; Founounou et al., 2013; Franz et al., 2018; Gho et al., 1999; Guirao et al., 2015; Lemke and Schnorrer, 2018; Levayer et al., 2015; Mauri et al., 2014; Michel and Dahmann, 2020; Prat-Rojo et al., 2020; Ray et al., 2015; Sarov et al., 2016). In particular, using spinning disc confocal microscopy, many studies have focused on the dynamics of a single tissue including the pupal wing and dorsal thorax (notum) tissues as well as histoblast nests. Yet, and as in most model systems, it remains critical to investigate multiple tissue and organ dynamics to define the underlying genetic and biophysical mechanisms coordinating the development of a whole organism. Here, we initially set out to test LSFM as an imaging approach to begin to explore genetic and mechanical coupling between multiple organs during *Drosophila* pupa development. While we could use LSFM to image non-embedded pupa, LSFM was not optimal for imaging cell and tissue dynamics with sufficient resolution. Consequently, we developed a Multi-View confocal microScope (MuViScope), which enables confocal multi-view imaging of non-immersed samples over time. We illustrate how its multi-view capability enables visualization of the dynamics of multiple organs, which will be instrumental to address fundamental biological questions related to organismal development, homeostasis and repair.

## Materials and Methods

### Confocal MuViScope microscope

A schematic outline of the MuViScope microscope (MuViScope) that was created using the 3D modeling software Solidworks (Dassault Systèmes SolidWorks Corporation, Waltham, MA, USA) is shown in Fig. 3. Technical drawings of the MuViScope and custom optical components are shown in Fig. S1. A comprehensive list of MuViScope components is also provided in Fig. S1B. The MuViScope consists of a multi-laser unit (GATACA systems, France), one spinning-disk confocal scan head (Yokogawa CSU-W1, Andor Technology) equipped with sCMOS (scientific complementary metal-oxide semiconductor) camera (Orca Flash 4.0 v2, Hamamatsu) and a home-made specimen holder, which is magnetically attached to a four-axis specimen-positioning system (x-y-z-θ). The system was built on an optical anti-vibration table (TMC, USA). The multi-laser bench (GATACA systems, France) is equipped with four laser lines 405 nm, 488 nm, 561 nm and 642 nm. The laser beam is guided through a single-mode fiber into the confocal spinning disk head. The optical path between the spinning head and the sample consists of two optical arms that direct light into two opposing dry objective lenses (Fig 3A, B, D). Each arm is composed of mirrors, a tube lens (TI-E 1×, Nikon Instruments Inc.) and an objective (CFI Plan Apo Lambda 10X/0.45 or CFI S Plan Fluor ELWD 40X/0.6, Nikon Instruments Inc.).

The collimated beam at the exit of the spinning-disk head is passing through a relay lens system and then diverted into the desired optical arm using a motorized flip mirror (custom-made by Errol Laser). The collected light follows the same optical path and is reflected from this flip mirror back onto the spinning disk head. A dichroic mirror and an internal detection-filter wheel located in the spinning disk head reflect the fluorescence to a sCMOS camera (Orca Flash 4.0 v2, Hamamatsu). The detection-filter wheel contains various detection filters (BrightLine long-pass and band-pass filters, Semrock) for imaging GFP and DsRed fluorophores. The capillaries (or rods) are inserted into a custom specimen holder made of stainless steel and held in place by the use of Teflon glands. The sample holder is then fitted into the axis of the rotating stage using a magnetic adapter. The whole assembly is held tightly and accurately to the four-axis specimen-positioning system (see Fig. 3 and Fig. S1). This four-axis (x-y-z-θ) specimen-positioning system consists of three motorized linear stages (2x M-404.1DG and 1xM-111.1DG, Physik Instrument) and one rotary stage (M-660.55, Physik Instrument) controlled by a 4-axes motion controller (C-884.4DC, Physik Instrument). With this system, the specimen holder can be translated along 3 axes and rotated around its main axis. A thermostatically controlled black plexiglass enclosure covers the whole unit. The temperature is controlled by the Cube temperature module (Life Imaging Services, Switzerland), and the humidity by a commercial humidifier (Eva, Stadler Form). These modules include sensors that are placed near the sample and maintain the user-defined temperature and humidity level. A cover at the top of the chamber provides access to the sample holder. A cold light source is available to help for sample positioning using brightfield illumination.

The optomechanical devices of the MuViScope are all operated through Metamorph software (Molecular Devices, version 7.8.13). The motorized flip mirror is controlled by a custom-made controller (Errol Laser, France) via a DAC (digital-to-analog controller).

### Drosophila stocks and crosses

Ecad:3xGFP (Pinheiro et al., 2017), *dpyOV1* (Bloomington #276), Mef2:GAL4>UAS-CD8:GFP (Mef2>CD8:GFP, gift from F. Schnorrer), hh-DsRed (hh-Pyr215, Kyoto DGGR 109137).

### Sample preparation

*Drosophila* pupae were staged by collecting white pupae (0 hours after puparium formation, hAPF). The puparium was removed for imaging except for the hh-DsRed experiment. Mounting and dissection was done under a binocular (Carl Zeiss). The posterior part of the pupal case was glued either (i) at the end of a capillary as done for other specimens in Laroche et al., 2019 using dental glue (Protemp II RF A3, 3M) for full rotation imaging (Fig. 3F); or (ii) the entire pupa was placed on double-sided tape (D6661933, 3M) on a flat platform at the end of a metal rod. The pupal case was then delicately dissected from the head to the beginning of the abdomen, either completely or only dorsal laterally. The capillary was then installed on its holder and inserted into the rotating stage. This mounting enabled unconstrained 360° rotation around the pupa’s anterior-posterior axis.

*Tribolium* transgenic *α*tub1-LifeAct:GFP, EF1*α*-nls:GFP (also known as LAN:GFP) (van Drongelen et al., 2018) adults were grown at 29°C in a Tupperware container containing flour and yeast. The *Tribolium* was euthanized by placing it at -20°C for 1h. It was then glued to the end of the capillary as detailed above. Imaging was carried out at 25 °C.

*Aranea* were collected in the wild in Paris (France) and were stored in ethanol before imaging. The Aranea was fixed to the end of the capillary by using dental glue. Imaging was done at 25 °C.

### Evaluation of pupa viability

Staged *Drosophila* pupae (30 hAPF) were lined up in a petri dish with the basal part on the plastic side. Four conditions were tested: pupae without glue, pupae glued on the ventral side with double-sided tape (3M), pupae glued at their end with dental glue (Protemp II RF A3, 3M), pupa glued at their end with UV activated glue (UHU, EAN:4026700481501). The petri dishes were placed in a box containing wet paper and placed in an incubator at 25°C for 7 days. The number of empty pupae was counted to assess the survival rate.

### Image Acquisition

The Metamorph acquisition software allows total control of the MuViScope via the Multi-acquisition mode. The control of the axes is done in the “Stage” tab of the “Multi Dimension Acquisition” menu (MDA) of the software. We use the Z2 stage function to control the rotating stage. The acquisition mode includes the acquisition of z-stacks of images, the creation of mosaics to allow the acquisition of the entire sample, the selection of multiple angles and the creation of time-lapses. All of these modalities can be combined without restriction. The exposure time per image was 300ms in binning 2 and a laser power of 0,6 mW at 488 nm and 1,5 mW at 561nm measured at the back focal plane of the objectives. For the z-stacks, the z-step was 4µm at the 10x objective and 2µm at the 40x objective with a step size from 100 and 500µm depending on the sample. The rotation speed of the stage was set at 360°/sec. The time between two acquisitions was either 15min or 30min.

The temperature of the thermostatically controlled chamber was set to 25°C with a humidity level of 70%. The sample was located using transmission light (white LED, Lumiled) and then finely focused using spinning disk confocal imaging by moving the motors in x,y,z, θ via the Metamorph software.

### Light-sheet fluorescence microscopy

Light-sheet microscopy was performed using the Lightsheet Z1 microscope (Carl Zeiss; illumination lens 10x/0.2 Carl Zeiss, – detection lens 20X/1.0 W Carl Zeiss) for the pupa immersed in PBS and the alpha3 light-sheet microscope system (PhaseView; illumination lens: EPIPLAN 10X/0.23 Carl Zeiss – detection lens MPLN 10X/0.25 Olympus) for the pupa w/o immersion. The mounting was identical as for the MuViScope (see § sample preparation).

### Image processing

Time-lapse images and multi-view z-stack acquisitions were processed using Fiji software (Schindelin et al., 2012). Briefly, for each angle and at each time point, a maximum projection (MIP) of z-stack images was made and then, if necessary, the resulting multi-view MIP images were stitched with the Pairwise Stitching plugin (Preibisch et al., 2009). This was done for every angle and every time point. A movie was then made from the sequential acquisitions and a montage showing the angles side by side was created by using the Stack Combiner plugin (Wayne S. Rasband, 1997). For Fig. 7, the images were denoised using ND-Safir software (Boulanger et al., 2010) and movie 5 was realigned using the ImageJ plugin StackReg (Thévenaz et al., 1998).

## Results

### Imaging the *Drosophila* pupa using LSFM

We set out to image multiple organs during *Drosophila* pupa development, focusing in particular on the wing and the dorsal thorax. In recent years, LSFM has become the reference for the *in toto* imaging of organisms (Wan et al., 2019). A known drawback of LSFM is the shadowing caused by absorption, scattering or refraction of light inside the sample (Rohrbach, 2009). Although these stripe artifacts can be found in any light microscope, they are more pronounced in light-sheet based microscopes due to the illumination from the side. Possible solutions to reduce shadowing include scanning illumination and/or multi-view imaging either by sample rotation or by dual-side sample illumination (Baumgart and Kubitscheck, 2012; de Medeiros et al., 2015; Huisken and Stainier, 2007; Keller et al., 2008). However, for samples of more than a couple hundred cells, these artifacts can undermine the ability to clearly image certain structures (Baumgart and Kubitscheck, 2012; de Medeiros et al., 2015; Huisken and Stainier, 2007; Keller et al., 2008). It remains unknown whether or not these artifacts might hinder the imaging of the *Drosophila* pupa. In addition, most commercial LSFM systems available require sample immersion in liquid or embedding in agarose. This potentially poses significant problems for organisms that rely on ventilation for gas exchange. Firstly, we opted for the light-sheet Z1 microscope from Carl Zeiss to assess the image quality using dual-illumination on the pupal dorsal thorax (Fig. 1A, B, B’). Upon immersion of the pupa in PBS, image quality was high on the lateral parts of the tissue and a cellular resolution could be obtained. However, even in this optimal imaging condition that precludes development due to pupa immersion, we did not obtain a high-quality image in medial regions of the tissue (Fig. 1B’). Secondly, we assessed LSFM image quality on the *Drosophila* pupa using the PhaseView ALPHA3 system (PhaseView, France). This system was chosen because its objective, detection lens and the light sheet alignment could be adapted to non-immersed samples. However, our tests indicated that the light sheet did not homogeneously illuminate the tissue when the dorsal thorax was imaged (Fig. 2A). Especially low signal was obtained in the medial part of the notum (Fig. 2A, insert). Likewise, when imaging the lateral side of the pupa the wing, legs and eyes were not homogeneously illuminated (Fig. 2B). In summary, these results illustrate that currently available commercial LSFM systems do not to produce high quality images in all areas of the dorsal thorax and wing tissues of the *Drosophila* pupa. This is likely due to the inherent shadowing artefacts associated with light-sheet microscopy.

**Figure 2.**
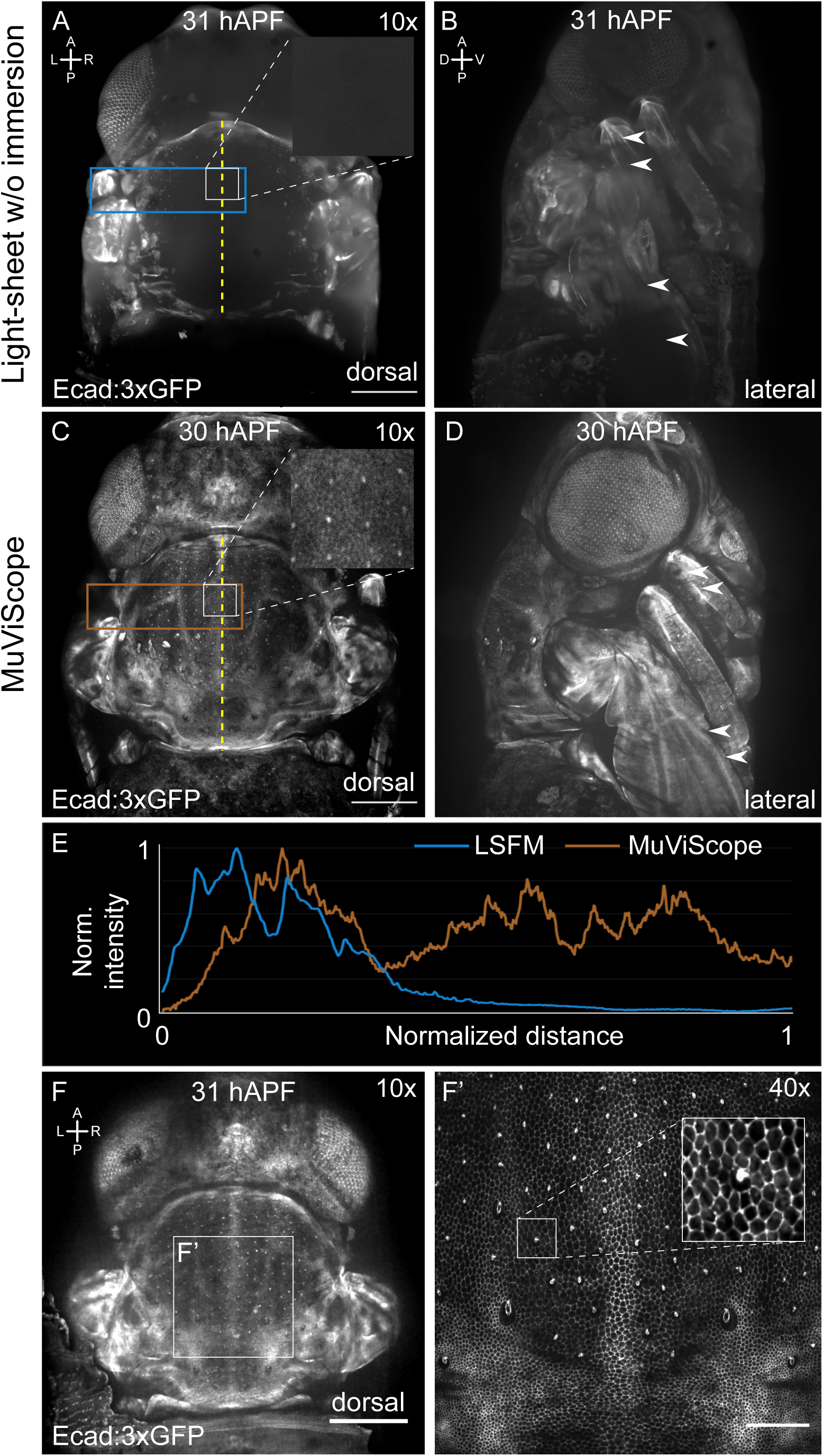
Comparison of light-sheet microscopy to MuViScopy. A-P and L-R axes in A for A,C and the A-P and Dorsal-Ventral (D-V) axis in B for B,D. Insert positions are indicated by white rectangle. A) Dorsal view of Ecad:3xGFP pupa without immersion imaged using the Phaseview ALPHA3 microscope at 10x adapted for non-immersed dual-side illumination light-sheet microscopy. Close-up in the square region (insert) where the image quality is lowest. The midline is indicated by a yellow dashed line. B) Lateral view of the pupa displayed in A. The arrowheads indicate positions where large intensity differences in the leg and wing that are due to shadowing effects (compare F). C) Dorsal view of Ecad:3xGFP pupa imaged using the MuViScope at 10x. Close-up in the square region (insert) to illustrate the image quality obtained in the medial region of the pupa. The midline is indicated by a yellow dashed line. D) Lateral view of the pupa shown in C. Arrowheads indicate regions approximatively corresponding to the ones indicated in B to illustrate the more homogeneous signal intensities observed in the leg and wing when imaging using the MuViScope. E) Graph of the normalized Ecad:3xGFP signal intensities within the rectangular regions shown in A (blue) and C (brown). Note that the normalized signal intensity in the sample imaged using LSFM (blue) goes to almost 0, while the MuViScope signal (brown) remains elevated throughout the sample. F) Dorsal view of Ecad:3xGFP pupa imaged with 10x and 40x objectives. The 10x view shows the overall animal shape. The white square region is imaged at higher resolution using the 40x objective and shown in F’. Cell outlines can be clearly ascertained in the insert in F’. Scale bars: 200 μm

### Imaging the Drosophila pupa using the MuViScope

Spinning disk confocal microscopy has often been employed to image individual epithelial tissues in *Drosophila* such as for example the wing, notum and the abdomen (Aigouy et al., 2010; Arata et al., 2017; Curran et al., 2017; Diaz-de-la-Loza et al., 2018; Etournay et al., 2015; Founounou et al., 2013; Gho et al., 1999; Guirao et al., 2015; Kanca et al., 2014; Keroles et al., 2014; Levayer et al., 2015; Michel and Dahmann, 2020; Prat-Rojo et al., 2020; Ray et al., 2015). Since individual tissues could successfully be imaged using confocal microscopy, we hypothesized that sample rotation along the anterior-posterior axis in combination with confocal microscopy could enable the visualization of multiple tissues. Furthermore, we foresaw that increasing the number of optical paths would enable imaging at multiple views at different magnifications. Accordingly, we developed the Multi-View confocal microScope (MuViScope) (Fig. 2A,C,E, Fig. 3, Fig. S1 and Movie 1).

**Figure 3.**
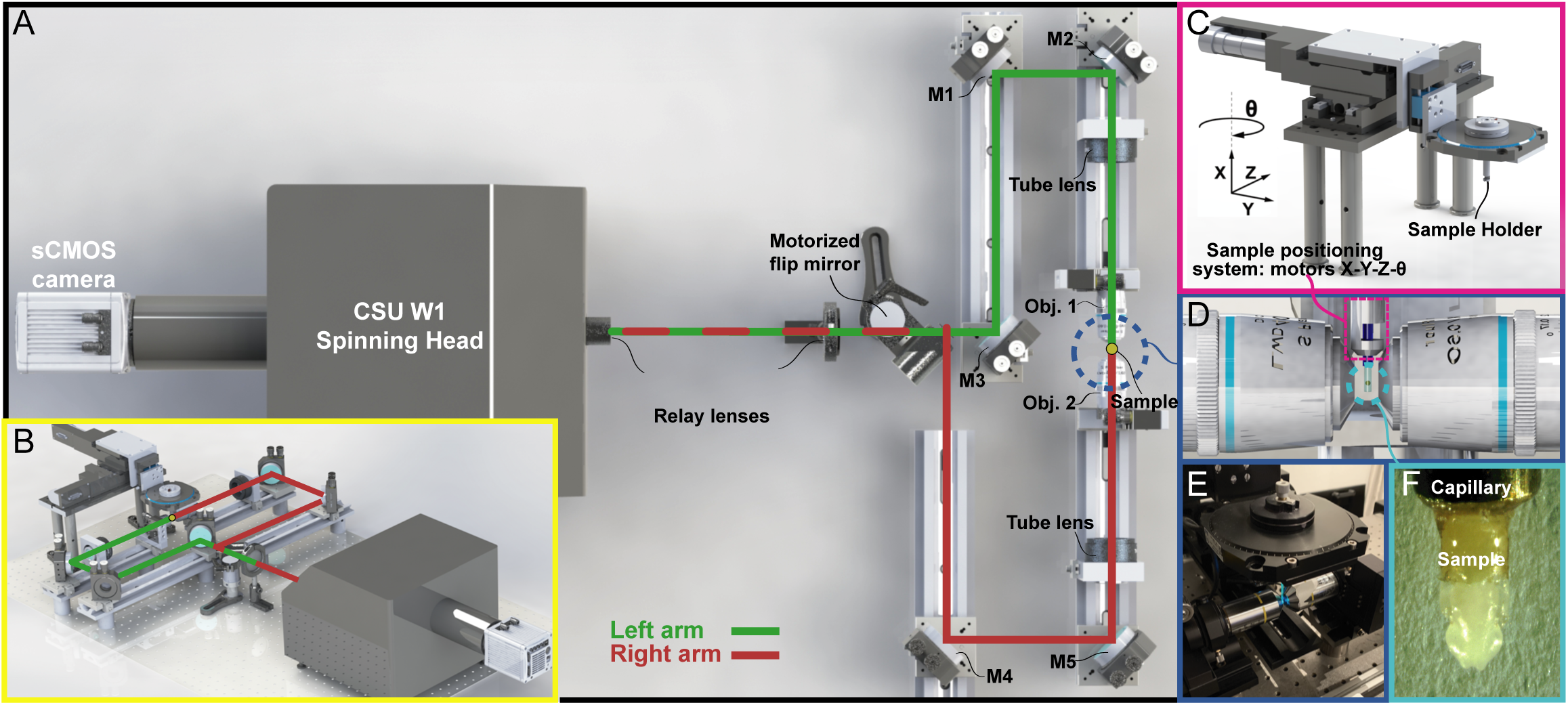
Multi-View confocal microScope setup (MuViScope). A) 3D rendering of the MuViScope setup. The MuViScope has two optical arms that illuminate the specimen and collect the fluorescence sequentially from the opposite side of the specimen: the optical pathway in red for the right arm and the one in green for the left arm. B) Perspective view of the microscope with the two-colored optical pathways. C) The capillary or the rod with the specimen is mounted in a custom-made sample holder. The specimen can be precisely positioned with 3 linear motors and 1 rotation motor to produce images from multiple views. D) The two opposing dry objective lenses are focused onto two different planes to image two opposite views of the specimen. E) Photo of the sample position system showing the sample holder and two objectives. F) The sample is maintained at the end of a capillary or on a metal rod.

The system is based on spinning disk illumination where the light source is composed of a fiber laser bench that is connected to the spinning disk head. The laser beam is focused on the sample by means of mirrors, dichroic mirrors, and lenses. The sample is imaged by using two dry long working distance objectives located on opposite sides of the sample. Each objective corresponds to an optical path where the excitation and collection of light is done via the same objective (Fig. 3A, B). The light emitted from the sample is collected by the excitation lens and passed back to the dichroic mirror located in the spinning disk head, which will reflect the light back to the sCMOS camera. The collection of the image via the opposite lens is done in the same way by changing the optical path using a motorized flip mirror (Fig. 3A, B, Movie 1). The sample is placed at the end of a capillary or metal rod which is held by a capillary holder (Fig. 3C-F). Sample mounting is done using dental glue on a capillary or double side tape on a small metal rod with a flattened end (Fig. 3F). The capillary and metal rod fit into a rotating stage that can be precision controlled along the x-y-z-θ axes (Fig. 3C-F, Fig. S1). Temperature and humidity are controlled within an opaque box, which surrounds the sample, sample holder, x-y-z-θ precision stages and objectives.

First, we assessed the ability of the MuViScope to visualize the dorsal thorax and wing tissues in *Drosophila* pupae. From the dorsal view, lateral structures such as the eyes were well imaged as well as the medial structures like the midline in the notum (Fig. 2C). The eye, legs, wing and wing veins could be visualized by rotating the pupa 90° (Fig. 2D). These results stand in contrast to the Phaseview microscope experiments, for which the notum and wing of living pupae could not be visualized without shadowing effects (Fig. 2A, C inserts, E). Furthermore, 180° sample rotation combined with 2 optical arms of the MuViscope offers the possibility to rapidly image with distinct objective magnifications to obtain both global and cellular tissue views in the same animal (500 ms apart, Fig. 2F,F’). The MuViScope therefore complements LSFM by permitting higher quality visualization of non-immersed samples.

### Whole animal imaging using the MuViScope

We next explored the capacity of the MuViScope to image samples using fluorescence and autofluorescence at the animal scale. We first imaged a pharate adult Ecad:3xGFP *Drosophila* pupa over 90° (Fig. 4A). The resulting images clearly showed major structures like the body segments and the legs as well as smaller ones like the folds in the wing, ommatidia in the eyes, and the bristles on the thorax (Fig. 4A). To test the MuViScope on other model organisms, multiple views of an adult *Tribolium* and *Aranea* were obtained. Using only autofluorescence, the legs, body and eyes could be ascertained in an *Aranea* (Fig. 4B). In the *α*tub1-LifeAct:GFP, EF1*α*-nls:GFP transgenic *Tribolium* adult the body segments, legs and antennae were clearly distinguishable (Fig. 4C and Movie 2). In addition, the resolution was high enough to discern small dimples present on the surface of the outer chitin layer (Fig. 4C’). The MuViScope is therefore capable of generating high quality 360° views in different species.

**Figure 4.**
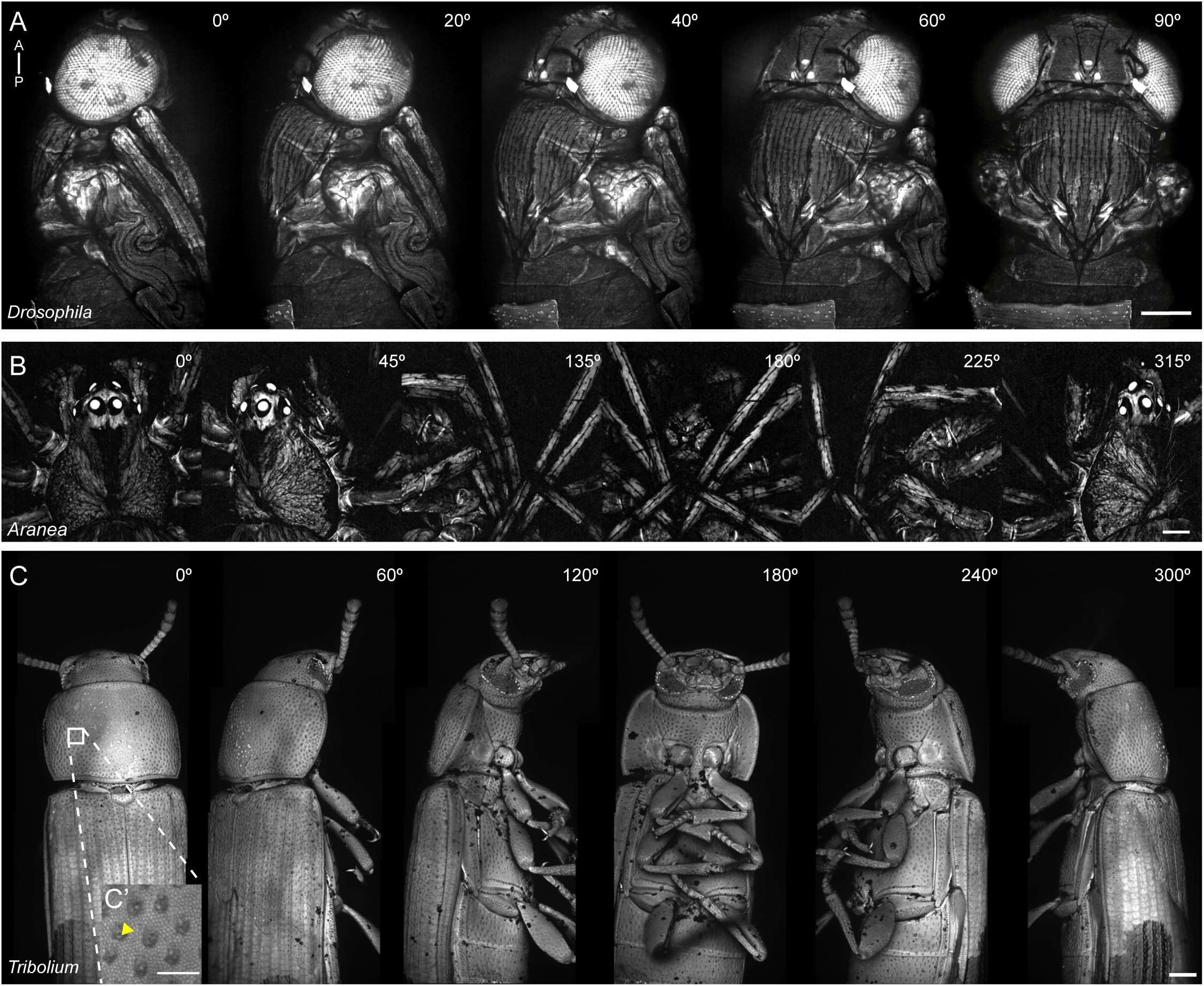
Application of MuViScopy to multiple model organisms. The A-P axis is indicated in A. Image collection is performed at 10x in A-C. A) 5-angle 90° view of an Ecad:3xGFP pharate adult *Drosophila*. B) 6-angle 315° view of a wild *Aranea*. Autofluorescence is used to image this sample. C) 6-angle 300° view of an *α*tub1-LifeAct:GFP, EF1*α*-nls:GFP adult *Tribolium*. See also Movie 2. C’) Close-up of the white box region in C. Yellow arrowhead: dimple in the outer chitin layer. Scale bars: A-C 200 μm, C’ 50 μm

### Imaging *Drosophila* pupa development to eclosion

Our next aim was to perform time-lapse live microscopy experiments. We first determined the optimal temperature and humidity for imaging. Viability was assessed by placing undissected pupae in the MuViScope imaging chamber with the temperature set to 25°C and humidity to 70%. Under these conditions high pupal viability was achieved (viability 90%, *n* = 9/10). Furthermore, pupal viability remained high upon removal of the pupal case (puparium) (viability 90%, *n* = 18/20), mounting with dental glue (viability 85%, *n* = 11/13) or on double-sided tape (viability 90%, *n* = 9/10). We then checked if pupae could be imaged from 0 hAPF to eclosion. For this, we used the transgenic hh-DsRed line, which is visible through the puparium and labels the posterior compartment of tissues (Akimoto et al., 2005). This line was selected to assess the MuViScope’s ability to image compartment domains during development. When performing 360° imaging using 4-angles (i.e. 90 degrees apart), Hh-DsRed signal could be observed on all sides of the pupa (Fig. 5A and Movie 3). We focused mainly on wing development. Labeling of the posterior compartment of the tissue allowed to observe that the wing posterior compartment undergoes a fast and global posterior movement at around 11 hAPF and extensive tissue twisting around 36 hAPF (Fig. 5A and Movie 3); two morphogenetic events occurring at the time of head eversion and wing folding respectively (Bainbridge and Bownes, 1981). These results show that the imaging conditions ensure high viability, and accordingly enables imaging of the entire pupal development.

**Figure 5.**
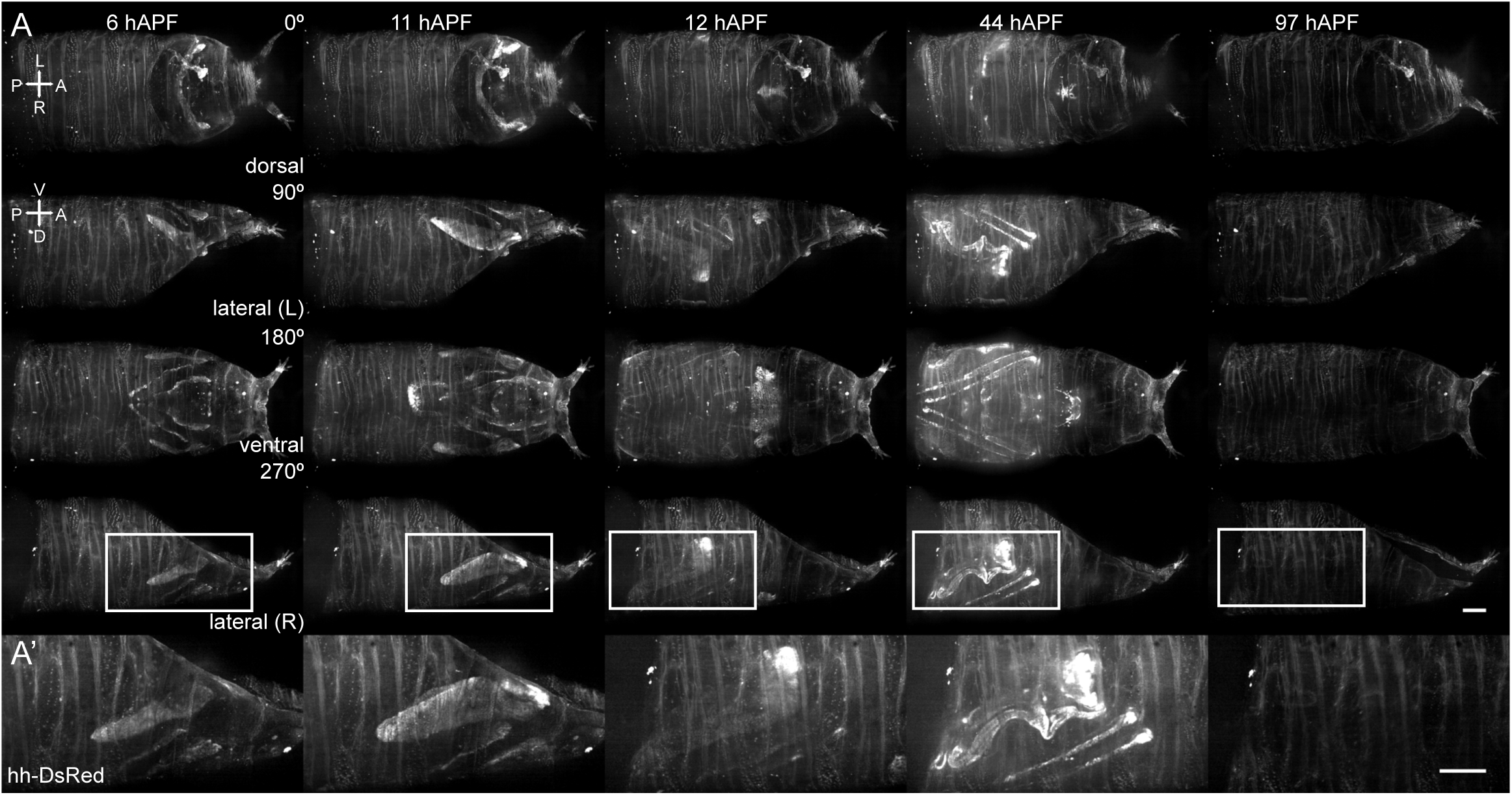
Imaging through the *Drosophila* pupal case using hh-DsRed throughout pupal development. A) 4-angle 360° view of a hh-DsRed pupa corresponding to dorsal, lateral right, ventral and lateral left views imaged at different time-points during pupal development. At time-point 97 hAPF the puparium is empty as the pupa has eclosed. The A-P and L-R axes are indicated on the dorsal view while the A-P and A-V are indicated on the lateral view. Image collection is performed at 10x. See also Movie 3. A’) Close-up of the lateral (R) view indicated by white boxes in A. Scale bars: 200 μm.

### Multiple muscle precursors imaging

Mesodermal tissues in the *Drosophila* pupa develop in multiple locations (Dutta et al., 2004; Jährling et al., 2010; Lemke and Schnorrer, 2018; McGurk et al., 2007). However, some mesodermal tissues in *Drosophila* have not been imaged live, because they develop inside the organism at positions for which mounting under a coverslip might be challenging. Especially difficult to image are the lateral and ventral muscles in the thorax, because the wing and legs respectively prevent placing a coverslip directly on top of the position where the muscles develop. 3-angle (0°, 90°, 180°) imaging was therefore performed to assess the performance of the MuViScope in imaging organs that develop internally. The striking pattern of dorsal-longitudinal muscles labelled by Mef2>CD8:GFP in the dorsal thorax could be well captured and imaged during development (Fig. 6, movie 4). An expansion in the anterior-posterior direction was visible between 26 hAPF and 50 hAPF, followed by a phase of muscle definition between 50 hAPF and 98 hAPF (Spletter et al., 2018). Moreover, sample rotation enabled the visualization of muscle development laterally and ventrally (Fig. 6, movie 4). Muscle expansion in the legs was most notable between 26 hAPF and 50 hAPF. This was followed by a refinement of the pattern of muscles from 50 hAPF and 98 hAPF similar to muscle development in the dorsal thorax. These results confirm and extend previous findings (Dutta et al., 2004; Jährling et al., 2010; McGurk et al., 2007; Spletter et al., 2018) and show that MuViScope is a suitable tool to explore the coordination of internal tissue development.

**Figure 6.**
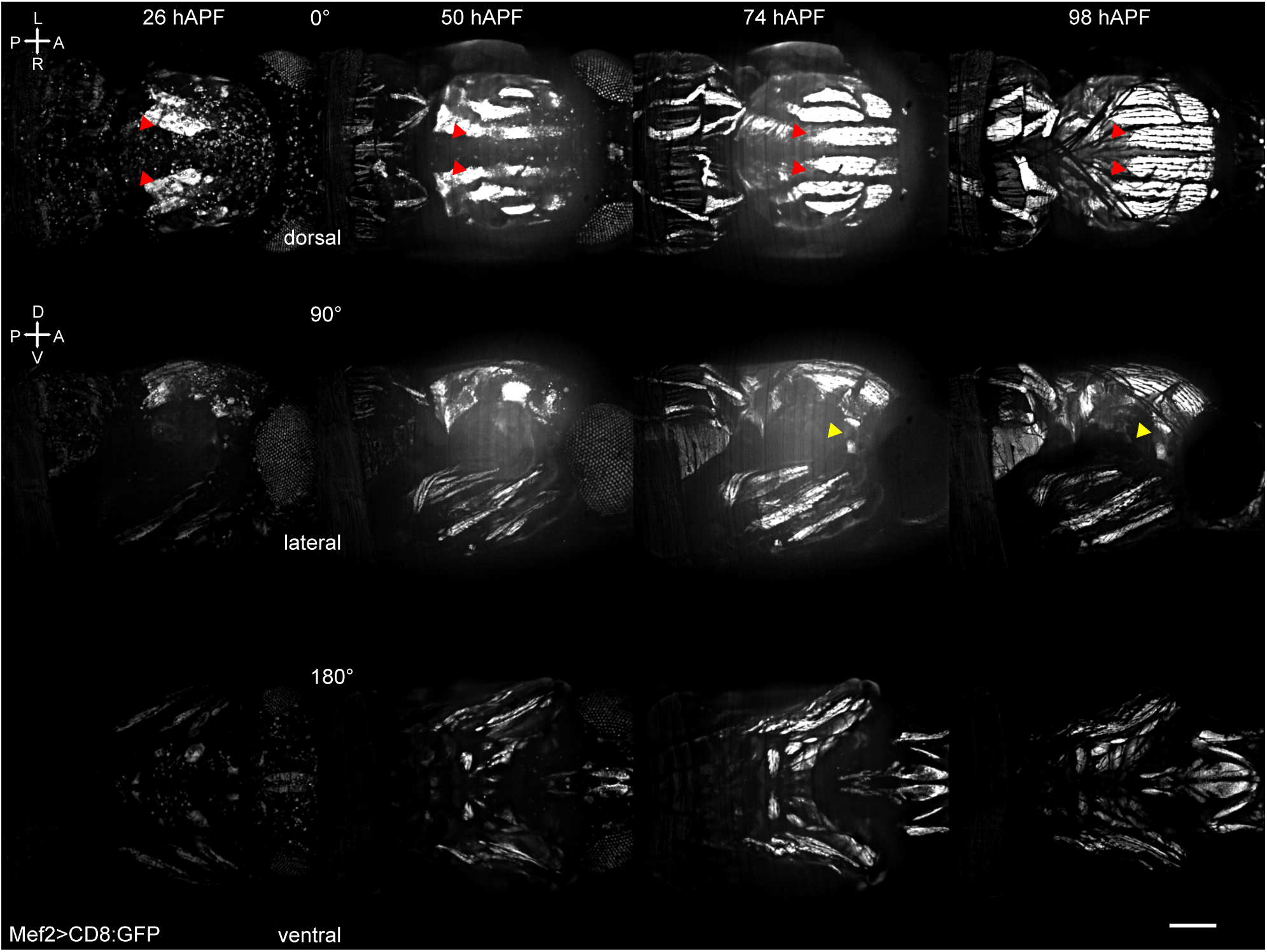
imaging of internal organs using the MuViScope. The A-P and L-R axes are indicated on the dorsal view while the A-P and D-V axes are indicated on the lateral view. 3-angle view of a Mef2>CD8:GFP pupa corresponding to a dorsal, lateral and ventral views imaged at different time-points during pupal development. Image collection is performed at 10x. dorsal-longitudinal muscles: red arrowheads; dorsal-ventral muscles: yellow arrowheads. See also movie 3. Scale bar: 200 μm

### Imaging multiple epithelial organs during development

In many cases, it might be advantageous to image multiple organs in the same animal to explore 1) whether and how genetic or mechanical developmental interplay exists between organs and 2) how systemic regulation coordinates animal scale development. We therefore set out to explore whether MuViScopy could be well suited to visualize multiple epithelial organs in the same animal. As a test case, we set out to characterize the development of both the wing and the dorsal thorax in both wild-type (wt) and *dumpy* (*dpy*) mutant conditions by labelling the tissue using Ecad:3xGFP. The extracellular matrix protein Dpy is known to control morphogenesis in the wing and dorsal thorax by enabling the attachment of the epidermis to the cuticle (Aigouy et al., 2010; Etournay et al., 2015; Metcalfe, 1970; Olguín et al., 2011; Ray et al., 2015). Previous research has shown that loss of Dpy function leads to misshaped wings and the formation of posterior lateral pits and anterior comma like invaginations in the notum (Aigouy et al., 2010; Etournay et al., 2015; Metcalfe, 1970; Olguín et al., 2011; Ray et al., 2015). Yet, the temporal sequence leading to formation of these defects in different organs has not yet been defined in the same animal using live microscopy. This therefore offers the opportunity to assess the ability of the MuViScope to explore tissue development in multiple organs.

We first visualized wing (lateral view) and dorsal thorax (dorsal view) tissues in Ecad:3xGFP over 90° to assess the wildtype processes at cell and tissue scale using both 10x and 40x imaging. In line with published results independently described in the wing and the notum (Aigouy et al., 2010; Etournay et al., 2015; Guirao et al., 2015; Ray et al., 2015), the lateral view allows to image the contraction the hinge starting at

∼17.5 hAPF and the subsequent elongation of the wing blade, whereas in the dorsal view we could observe the flows of thorax cell towards the anterior as well as the morphogenesis of the posterior lateral region of the notum starting at ∼20 hAPF (Fig. 7A,B, Movie 5). We then performed similar movies in *dpyOV1* animals. In *dpyOV1* prior to 17 hAPF before hinge contraction, the distal end of the wing and the dorsal thorax were similarly shaped compared to wt (Fig. 7A-D, green arrowheads). The distal part of the wing was round and no invaginations could be observed in the notum. Starting from ∼17.5 hAPF when the hinge started to contract, the distal part of the *dpyOV1* wing started to change its shape while moving towards the hinge. Accordingly, the wing blade failed to elongate as observed in wt tissues (Movie 5, Fig. 7A-D). Using both the lateral and dorsal view we could observe that similar to the wt tissue, the hinge contraction was followed in the notum by cell flow towards the anterior at ∼20 hAPF (Movie 5). However, in *dpyOV1* conditions the anterior comma like invaginations started to appear at ∼24 hAPF and became progressively more visible until 34 hAPF (Movie 5, Fig. 7C, yellow arrowheads). The posterior lateral pits in the notum appeared between 24 and 34 hAPF (Fig. 7C, red arrowheads). These pits could not be observed in wt tissue (Fig. 7A, yellow and red arrowheads). Together, these results illustrate that the MuViScope can be used to investigate the morphogenetic processes in both wt and mutant animals in multiple organs within the same animal.

**Figure 7.**
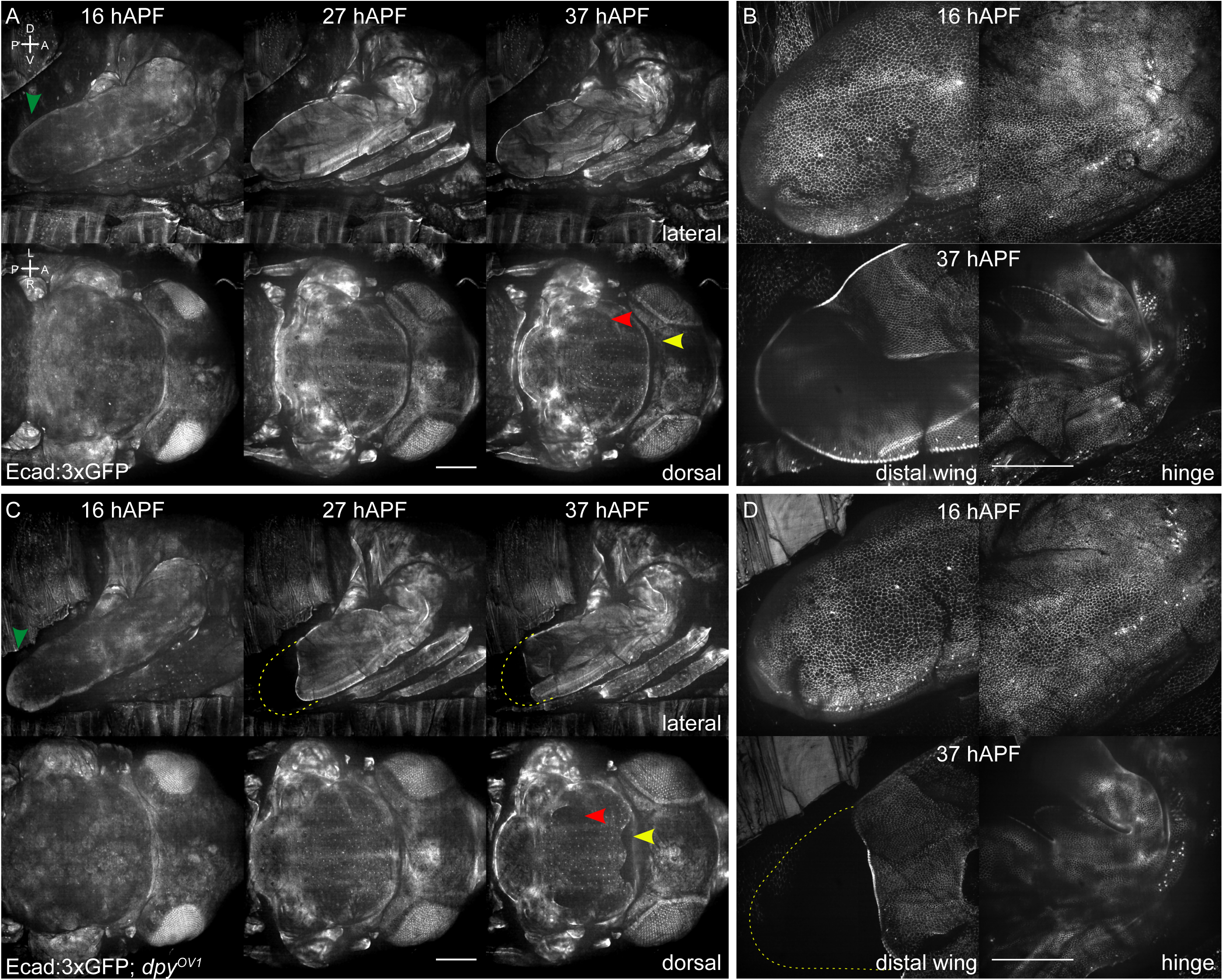
imaging of multiple organs in wt and *dpyOV1 Drosophila* pupae. Green arrowheads: time-point when distal wing is similarly shaped in both conditions. Yellow arrowheads: position of the anterior comma like invaginations in Ecad:3xGFP, *dpyOV1*. The arrowhead points to approximately the same position in Ecad:3xGFP, wt for comparison. Red arrowheads: position of a lateral pit in Ecad:3xGFP, *dpyOV1*. The arrowhead points to approximately the same position in Ecad:3xGFP, wt for comparison. A) 2-angle 90° views of Ecad:3xGFP pupa corresponding to the lateral and dorsal views at 16, 27 and 37 hAPF using a 10x objective on the first MuViScope arm. The A-P and L-R axes are indicated on the dorsal view while the A-P and D-V axes are indicated on the lateral view. B) Lateral view of the pupa shown in A using the 40x objective on the second microscope optical arm at two positions along the A-P axis to image both the wing most distal part (left) and the hinge (right). C) 2-angle 90° views of Ecad:3xGFP, *dpyOV1* pupa corresponding to the lateral and dorsal views at 16, 27 and 37 hAPF using a 10x objective on the first MuViScope arm. Dashed lines: delineates approximate shape of a wt wing. D) Lateral view of the *dpyOV1* pupa shown in C using the 40x objective on the second microscope arm at two positions along the A-P axis to image both the wing most distal part (left) and the hinge (right). Dashed lines: delineates approximate shape of a wt wing. Scale bars: A,C 100µm and B,D 50µm. See also movie 5

## Discussion

Here, we implemented an advanced microscopy methodology named MuViScopy based on spinning disk confocal illumination and multidirectional imaging by sample rotation and double-sided illumination/detection. We showed that this methodology can be used to image Arthropods such as *Tribolium*, *Aranea* and *Drosophila*. In particular, we explored the ability to visualize organs and multiple organ development in *Drosophila*. MuViScopy enables high quality automated 360-degree imaging of living biological samples without sample embedding/emersion. It’s conventional sample illumination and fluorescence collection by a single objective offers clear advantages compared to light-sheet imaging. It enables homogeneous sample illumination without the need to post-acquisition image processing to reconstruct one homogeneously illuminated field of view. External organs such as the epidermis were well captured throughout the entire thorax and could be visualized with cellular resolution. Furthermore, MuViScopy permitted live imaging of the morphogenesis of internal organs that are challenging to image due to their location in the organism. In addition, multidirectional imaging enables to assess how the relative developments of distinct organs unfold in both wt and mutant animals with distinct objective magnifications on the same sample. MuViScopy therefore opens up a new range of experimental opportunities to better understand how multiple organs develop as well as to explore the genetic, biophysical and systemic regulations that underlie the organ dynamics. We therefore believe that the MuViScope will be advantageous for the study of fundamental biological questions in organisms that rely on ventilation for gas-exchange. Accordingly, we have provided the MuViScope blueprint and detailed technical information to support the dissemination of MuViScope microscopy.

We see room to improve the MuViScope in multiple ways. First, a sample finder module as is present on the commercial Carl Zeiss Z1 system could reduce the time spent finding the optimal focus. Second, the design of the sample holder was inspired by light-sheet microscopy. This entails that the sample holder currently only fits one pupa. Enlarging the mounting area would increase sample throughput at the cost of reducing the number of angle position that can be imaged. Third, photon collection could be optimized by using a higher quantum efficiency sCMOS camera and a field homogenizer module. Lastly, the addition of new electrically tunable lenses would increase the acquisition speed of the z-stacks by enabling successive z planes imaging without moving the sample. We foresee that these future changes will further enhance the range of applications of the MuViScope in the study of fundamental biological questions.

## Acknowledgments

We would like to thank: Frank Schnorrer (Institute for Developmental Biology Marseille) and Pierre Leopold (Institut Curie) for *Drosophila* stocks, Maurijn van der Zee and Kees Koops (both Leiden University) for *Tribolium* stocks, France Lam (Institute of Biology Paris-Seine) for access to and help with the Phaseview Alpha3 system, the cellular imaging platform at SFR Necker for access to the Zeiss Z1 lightsheet microscope and Sébastien Dupichaud for his help in operating it, Maité Coppey (Institut Jacques Monod) for help with writing funding applications, Christine Rollard (Muséum national d’Histoire naturelle) for help with spider identification, François Graner and Isabelle Bonnet for the fruitful discussions and preliminary tests at the beginning of the project and Aude Maugarny-Calès for her advice and help with *Drosophila* sample mounting. The MuViScope was financed by France-BioImaging national research infrastructure (ANR-10-INBS-04) and additional funding from ERC Advanced (TiMorph, 340784), ARC (SL220130607097), ANR Migrafolds (ANR-18-CE13-0021), ANR Labex DEEP (11-LBX-0044, PSL ANR-10-IDEX-0001-02) grants. EvL was financed by the ITN ’PolarNet’ (675407), EvL and AV were both financed by the FRM (FDT201904008163, FDT201805005805).

## Movie legends

**Movie 1 | Animation of the MuviScope 3D blueprint.** Schematic illustration of the different components of the MuViScope. The movie first shows a 3D top view plan of the system with on the right side the spinning disk head and the camera, motorized flip mirror and the two paths (right and left arms) on the central part and the specimen-positioning system on the left side. The animation then shows a side view with a full rotation followed by a zoom on the motorized flipping mirror, the two objectives and the specimen holder with the positioning system. The plan is drawn to scale.

**Movie 2** | **Movie of a full 360° rotation of an adult *Tribolium*.** The A-P axis is indicated. Large structures like body segments can be observed, as well as smaller dimples on the body. The *α*tub1-LifeAct:GFP, EF1*α*-nls:GFP (LAN:GFP) transgenic *Tribolium* line was used for the experiment. Scale bar 200 μm.

**Movie 3** | **Time-lapse imaging of the posterior compartment using hh-DsRed during Drosophila pupa development.** The A-P and L-R axes are indicated on the dorsal view while the A-P and D-V axes are indicated on the lateral view. Imaging is started at white pupa (0 hAPF). A quick posterior movement occurs between 10.5 and 11 hAPF. Extensive tissue shape changes take place between 35 and 45 hAPF. The fly ecloses at 96 hAPF. Scale bar 200 μm.

**Movie 4 | Time-lapse imaging of muscle development using Mef2>CD8:GFP.** The A-P and L-R axes are indicated on the dorsal view while the A-P and D-V axes are indicated on the lateral view. Dorsal longitudinal muscles compact between 26 and 34 hAPF, before elongating in the anterior posterior direction. The muscles span the entire dorsal thorax at 46 hAPF. In the lateral and ventral view muscle development the signal in the muscles in the legs increases between 26 and 54 hAPF before plateauing. In the lateral view a small dorsal-ventral muscle can be seen to appear at 58 hAPF. Scale bar 200 μm.

**Movie 5** | **Time-lapse imaging of Ecad:3xGFP, wt and *dpyOV1* pupae**. The A-P and L-R axes are indicated on the dorsal view while the A-P and D-V axes are indicated on the lateral view. Organ imaging in Ecad:3xGFP, wt and Ecad:3xGFP, *dpyOV1* pupa. Points of interest are indicated with arrows. Green arrowheads: time-point when distal wing is similarly shaped in both conditions. Yellow arrowheads: position of the anterior comma like invaginations in Ecad:3xGFP, *dpyOV1*. The arrowhead points to approximately the same position in Ecad:3xGFP, wt for comparison. Red arrowheads: position of a lateral pit in Ecad:3xGFP, *dpyOV1*. The arrowhead points to approximately the same position in Ecad:3xGFP, wt for comparison. Scale bars 50 μm at 40x and 100 μm at 10x.

---

**Supplementary Figure 1.**
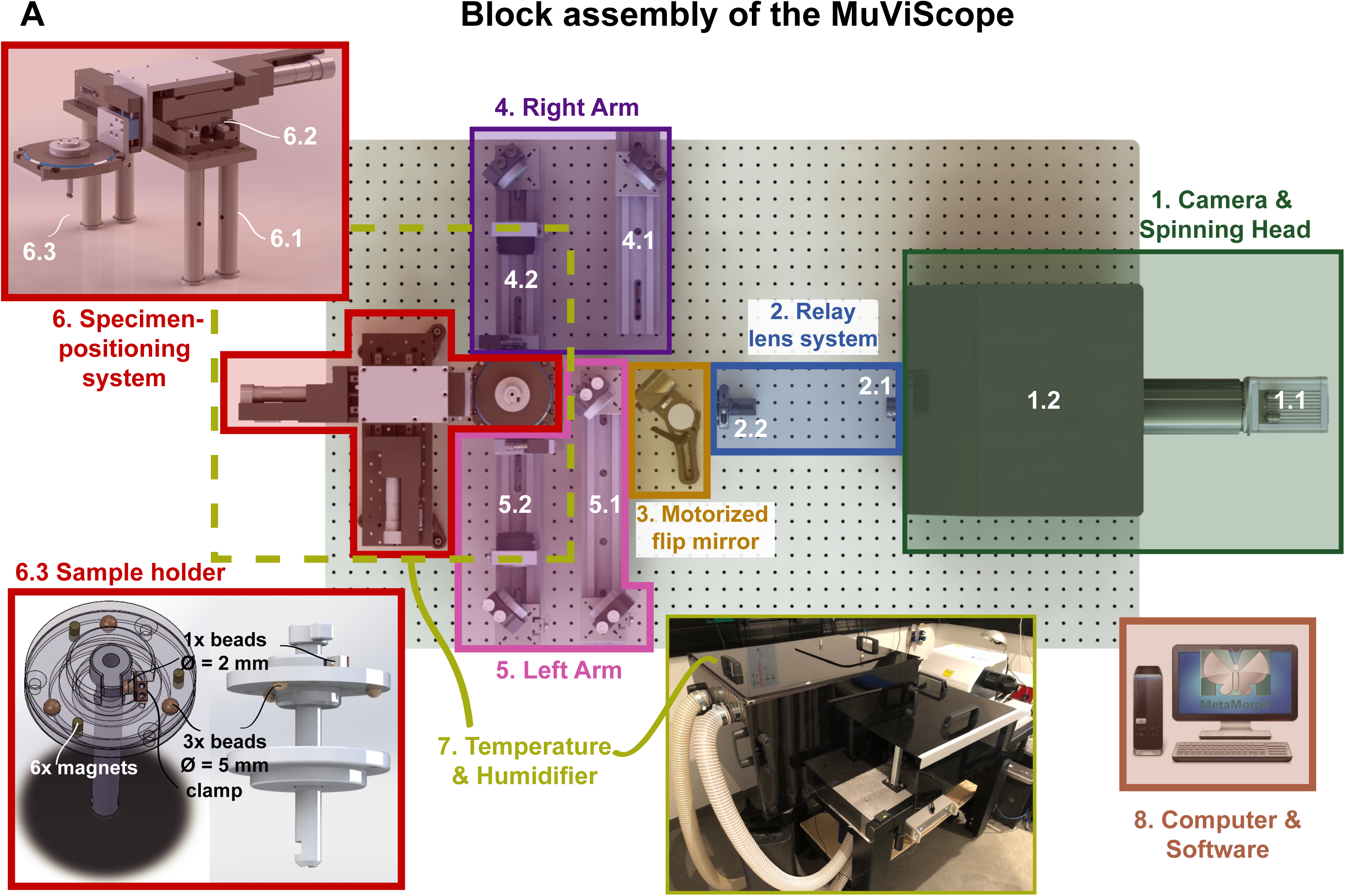
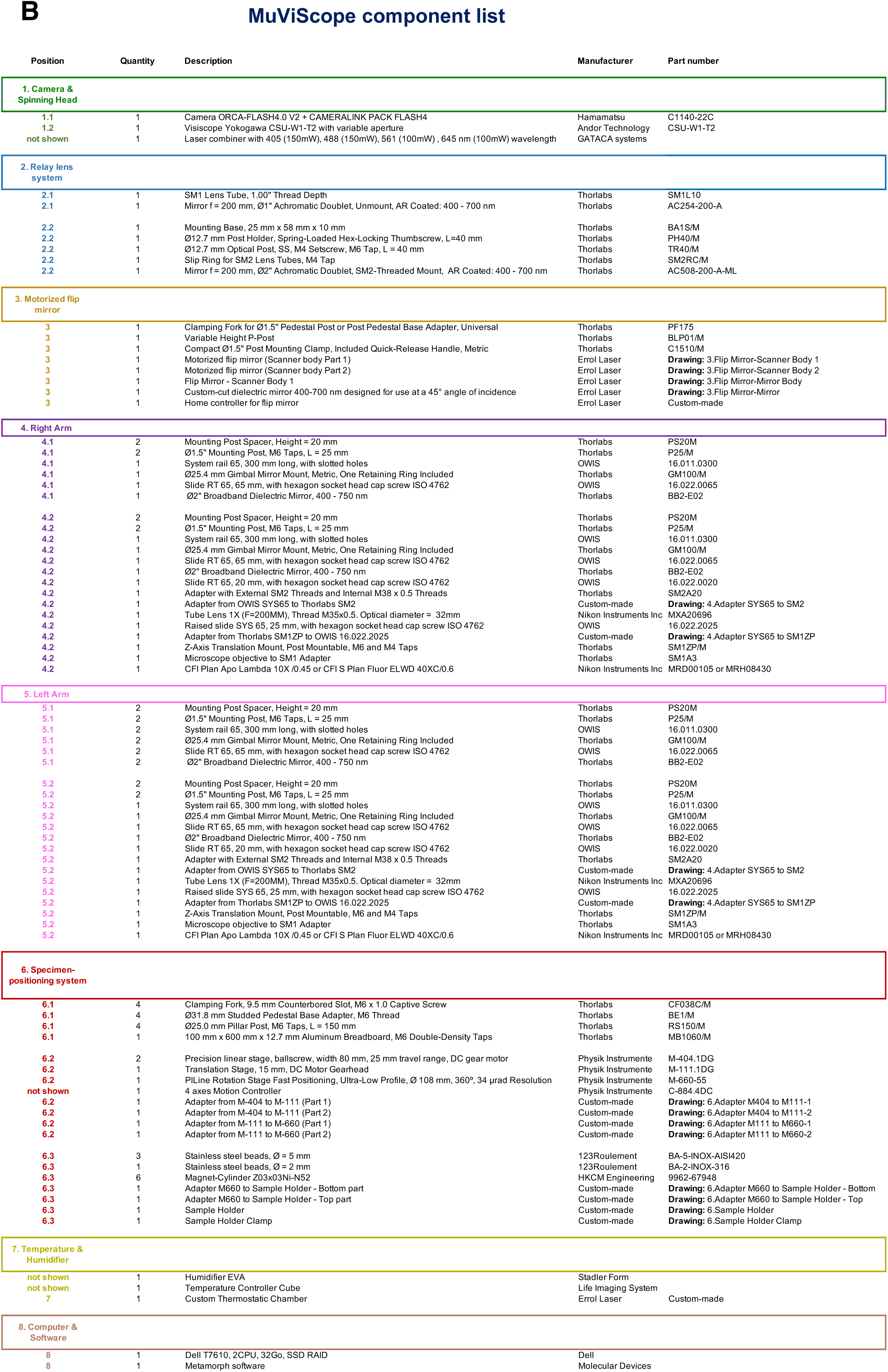
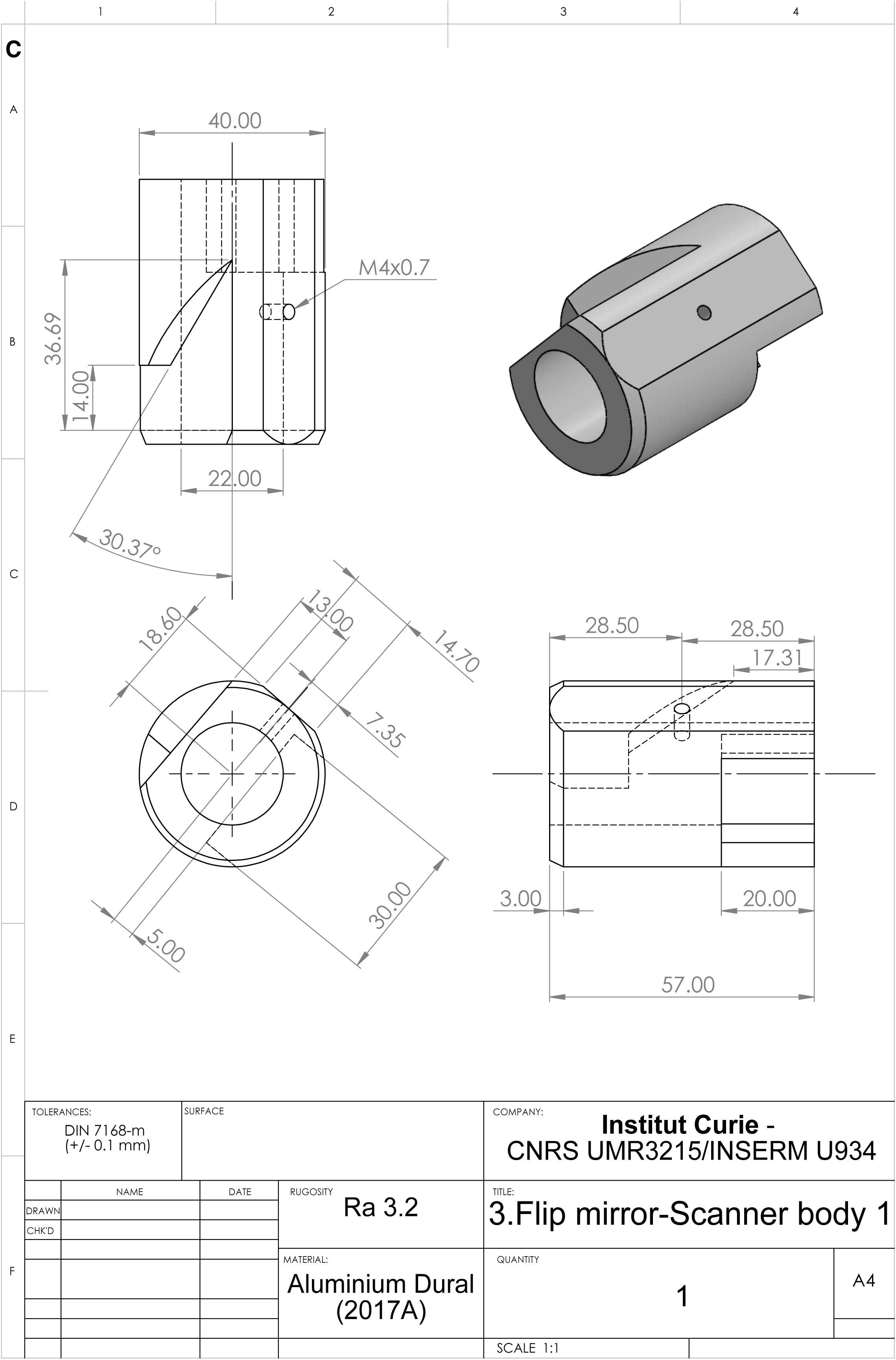
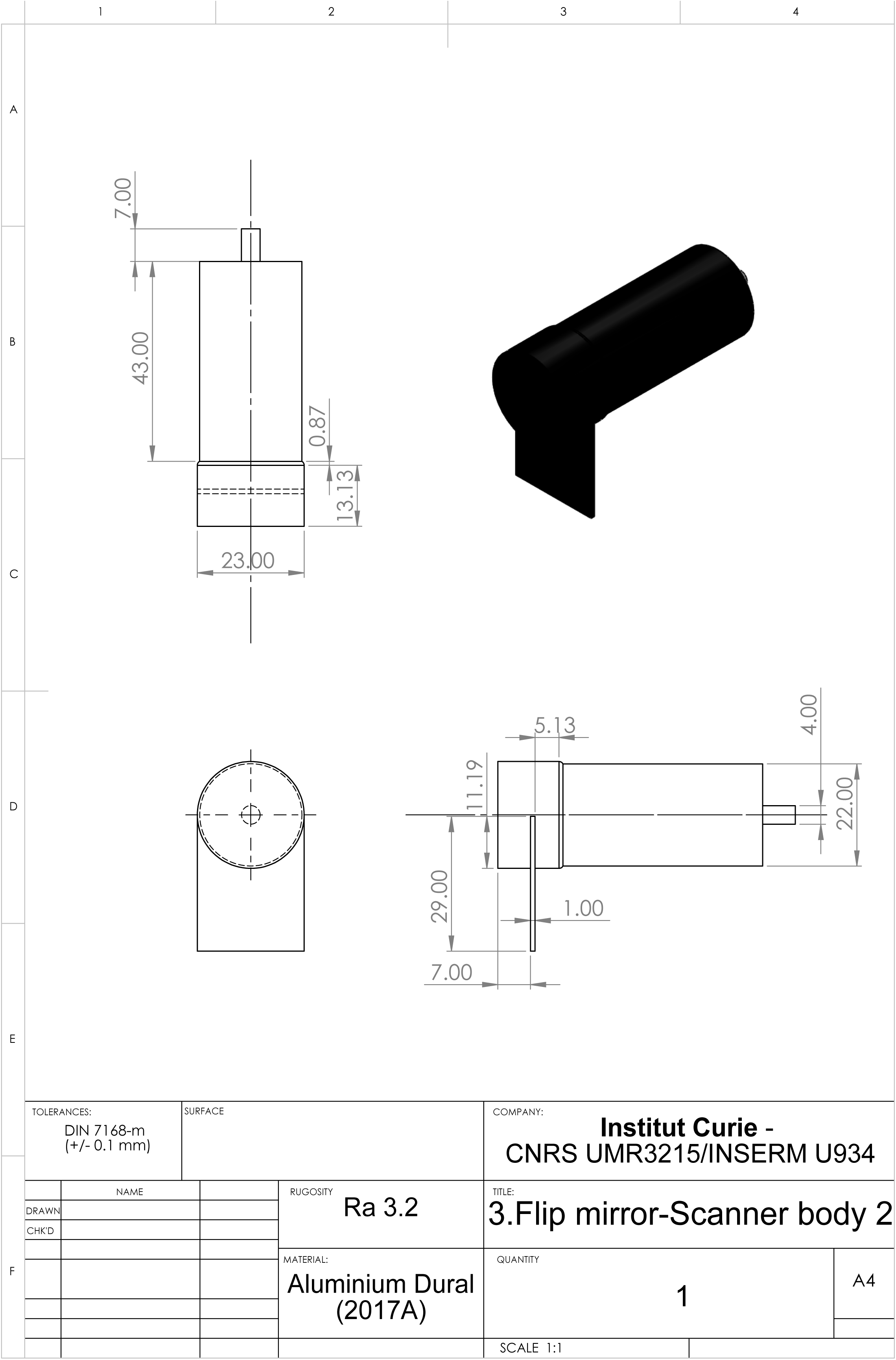
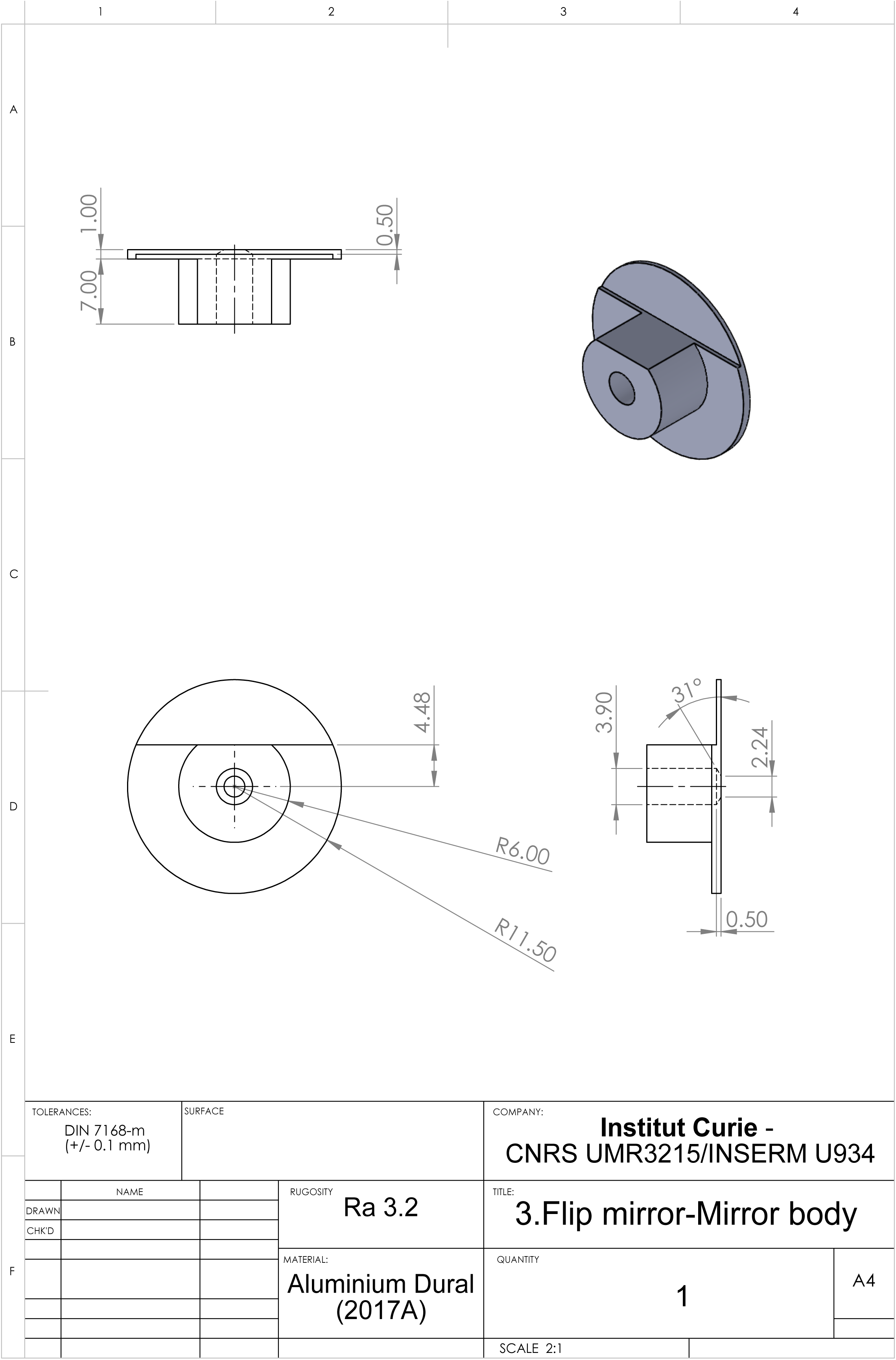
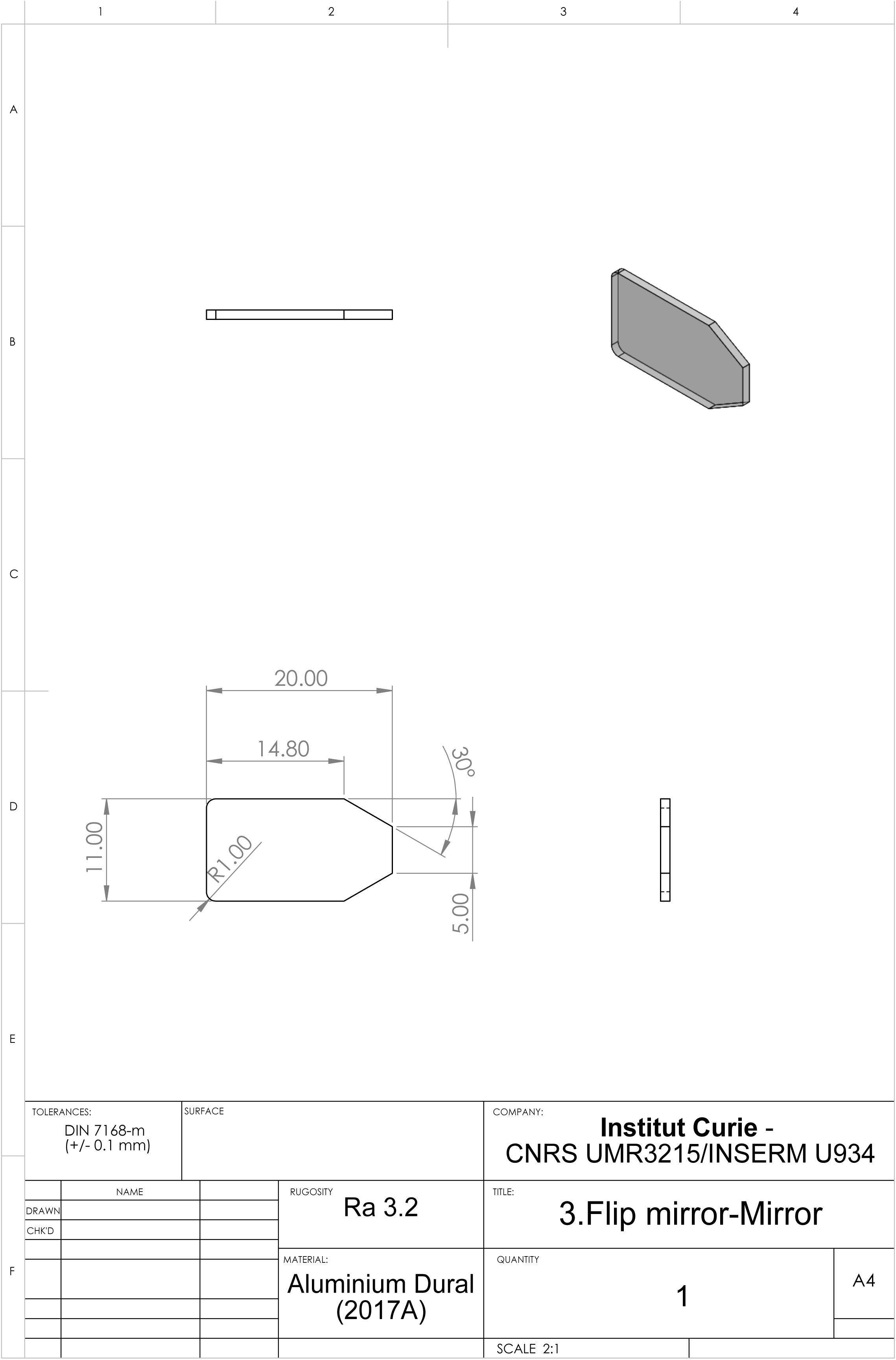
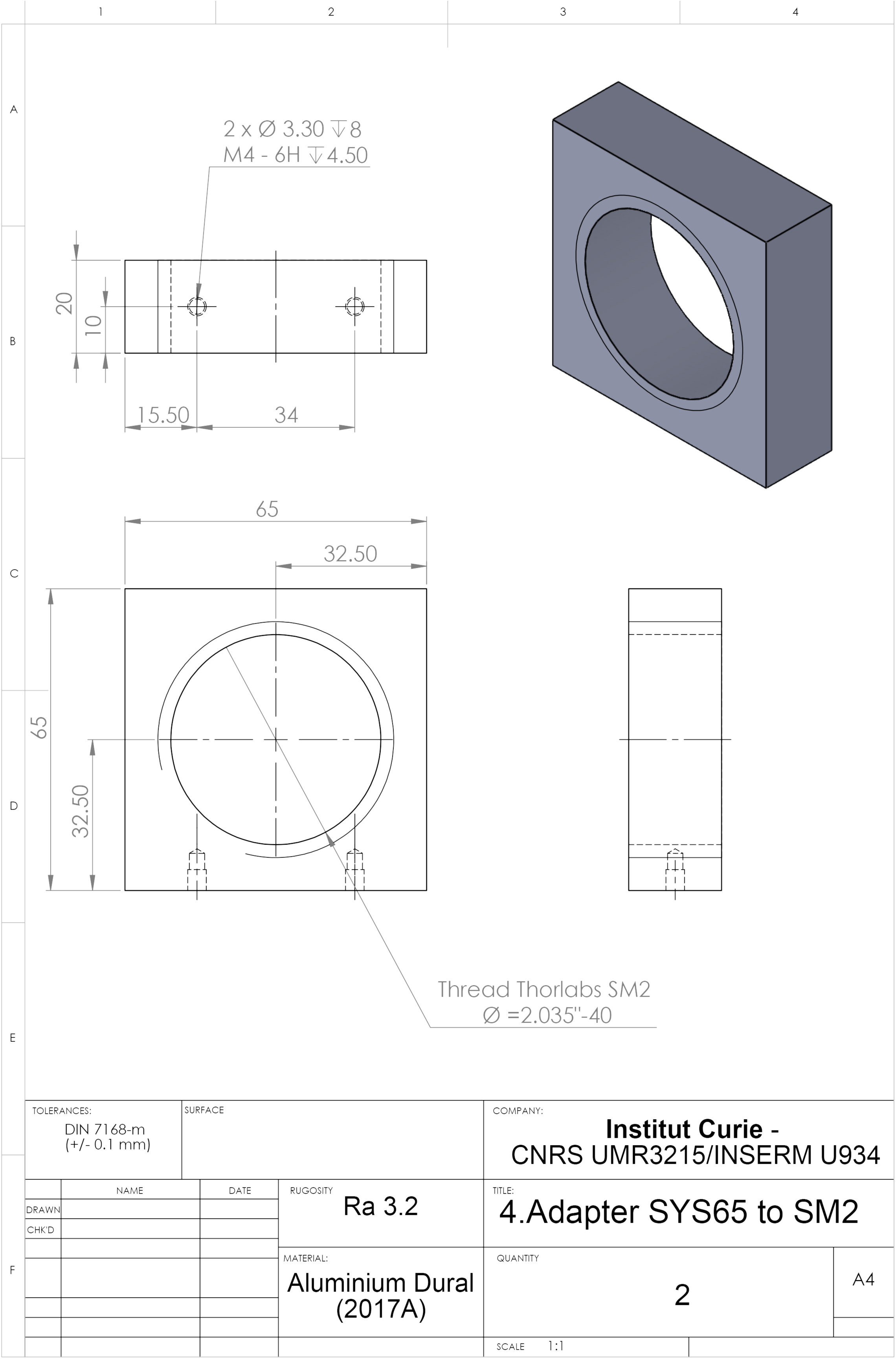
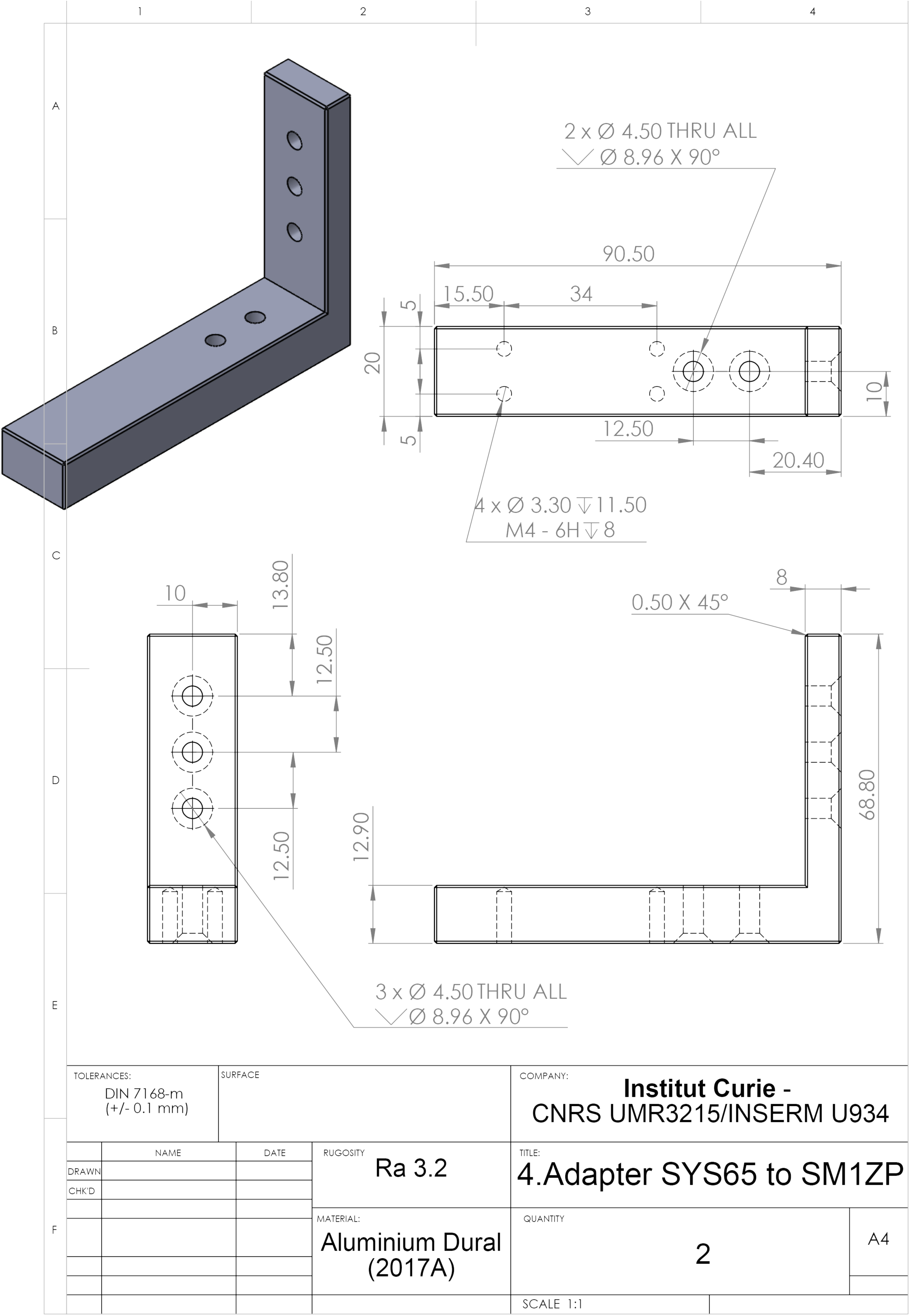
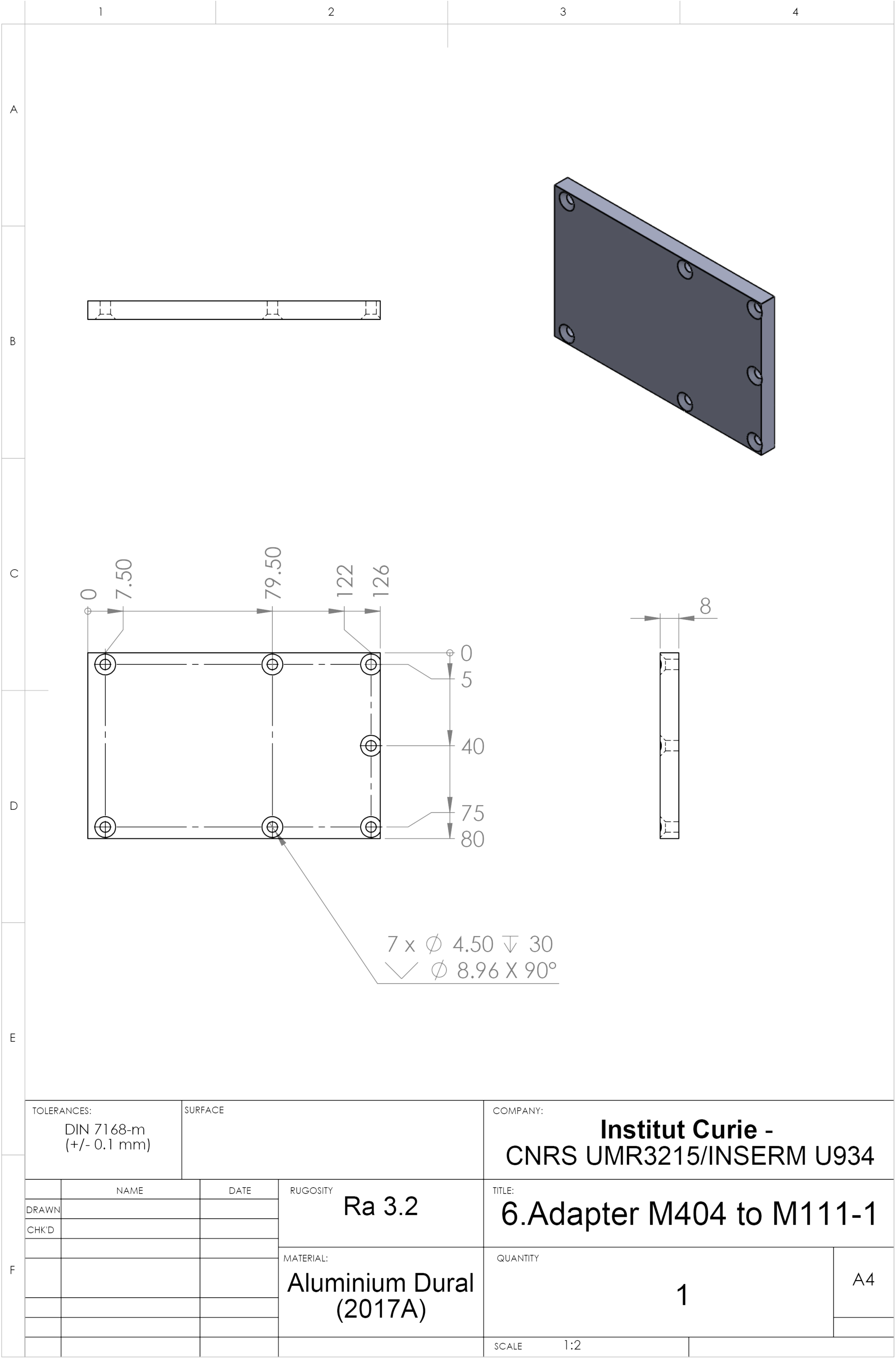
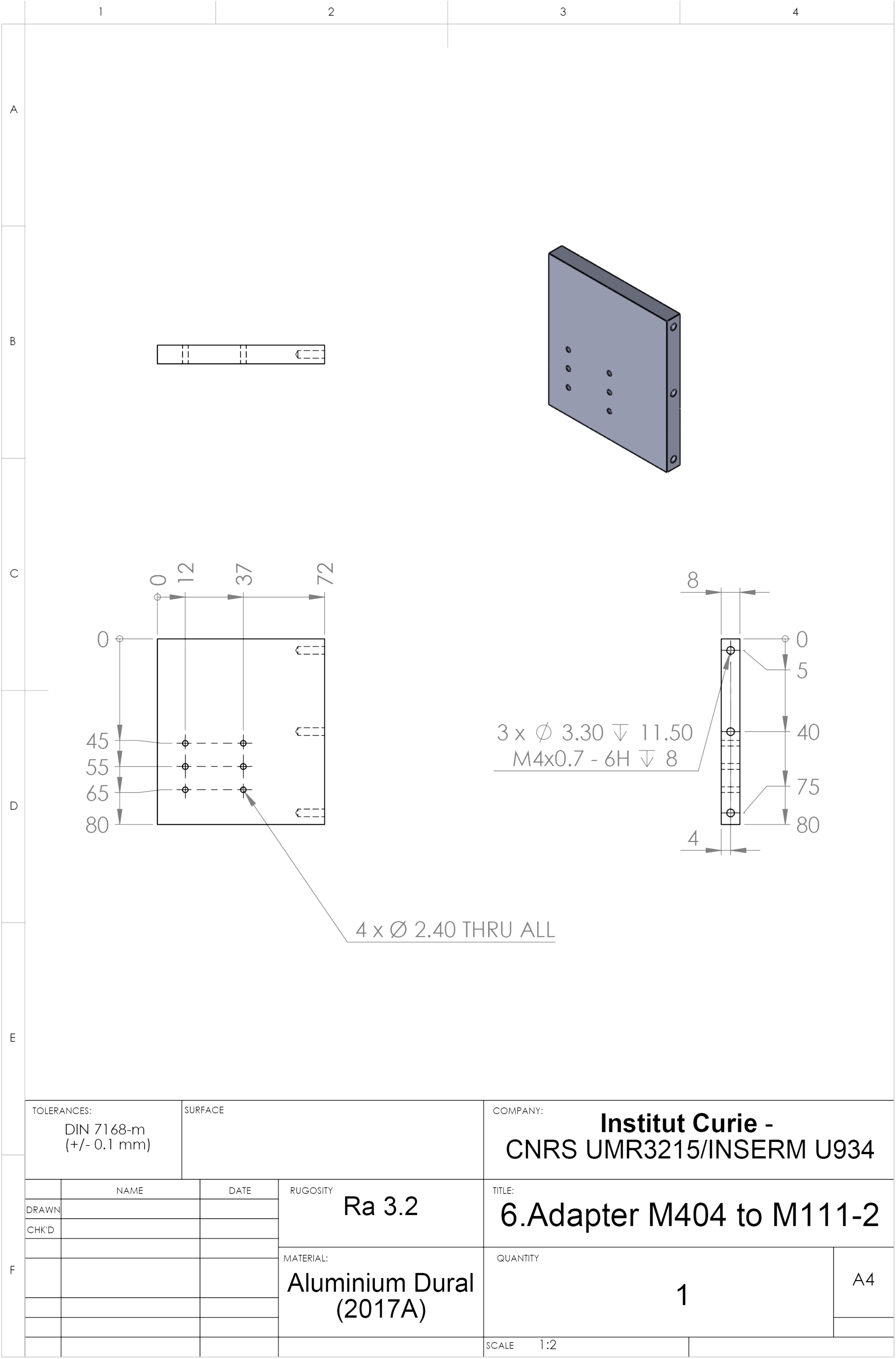
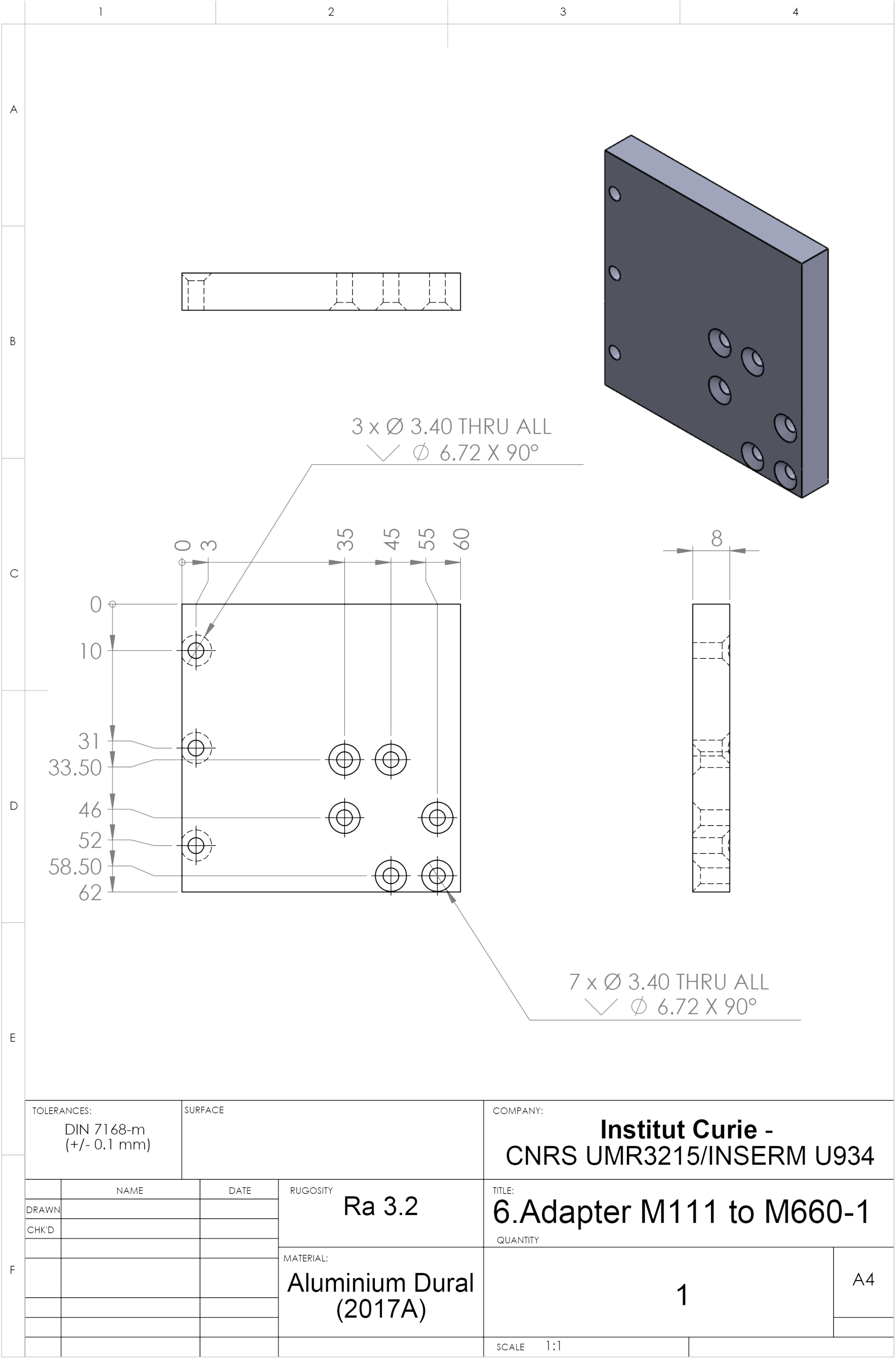
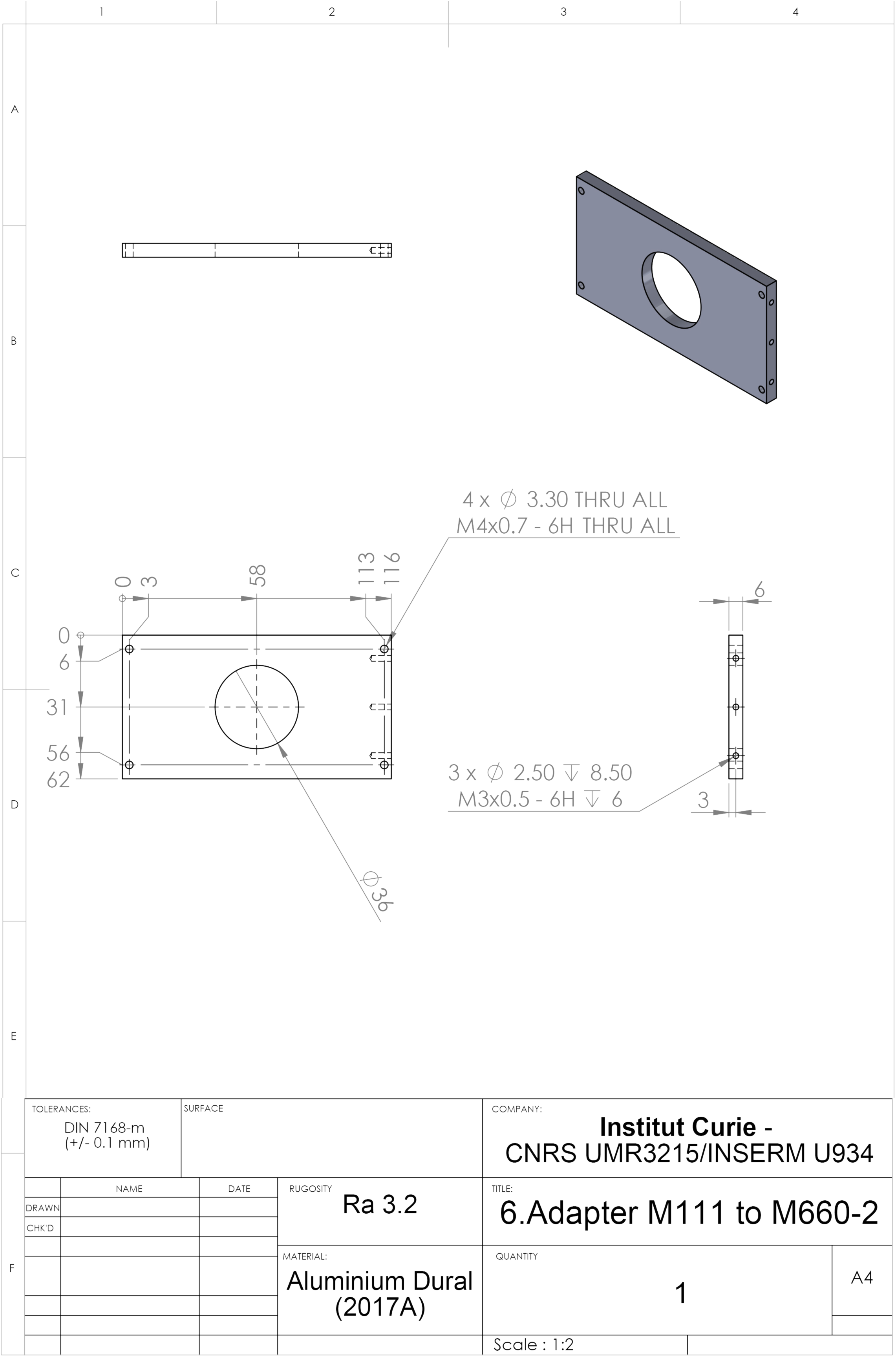
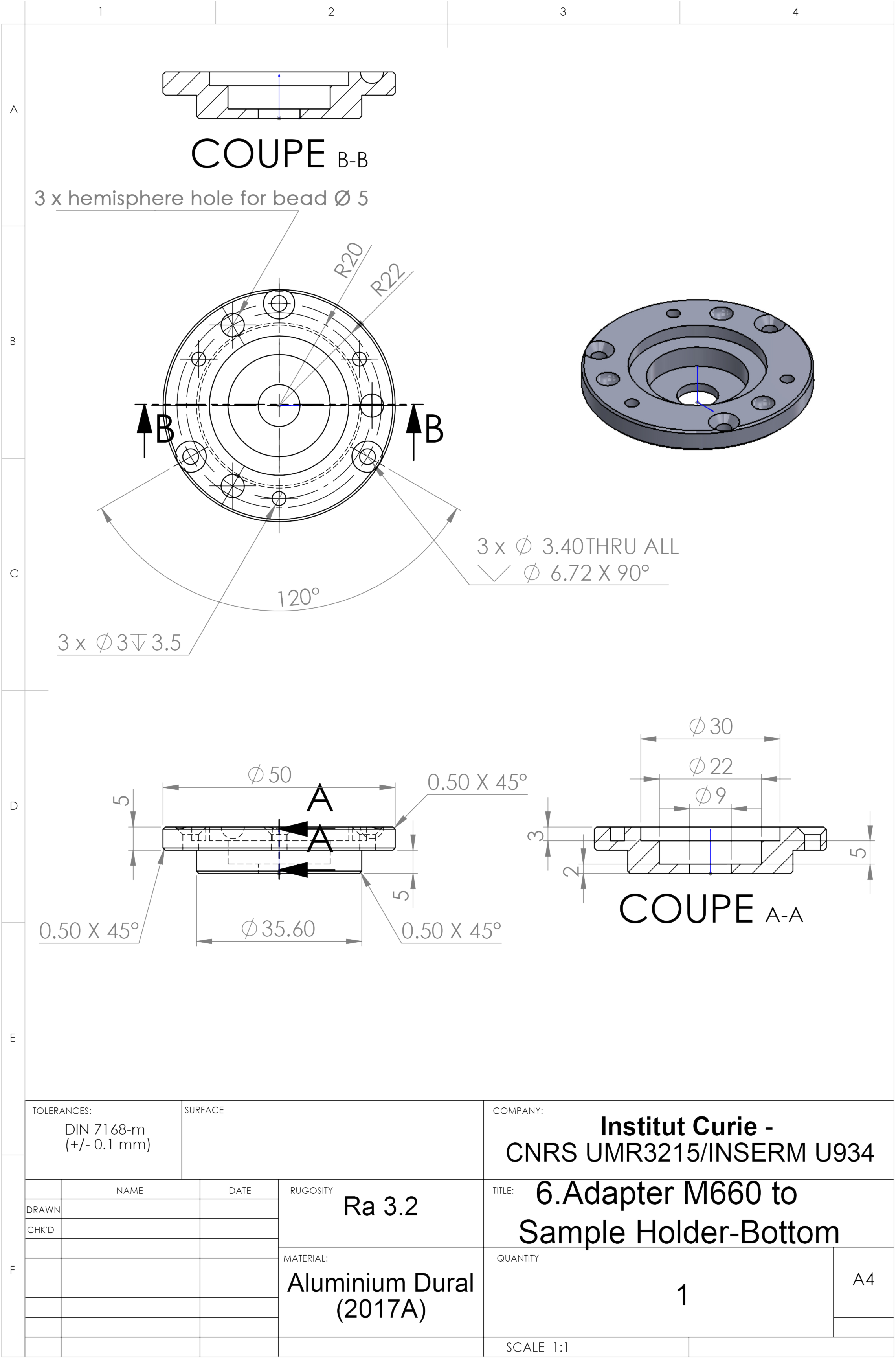
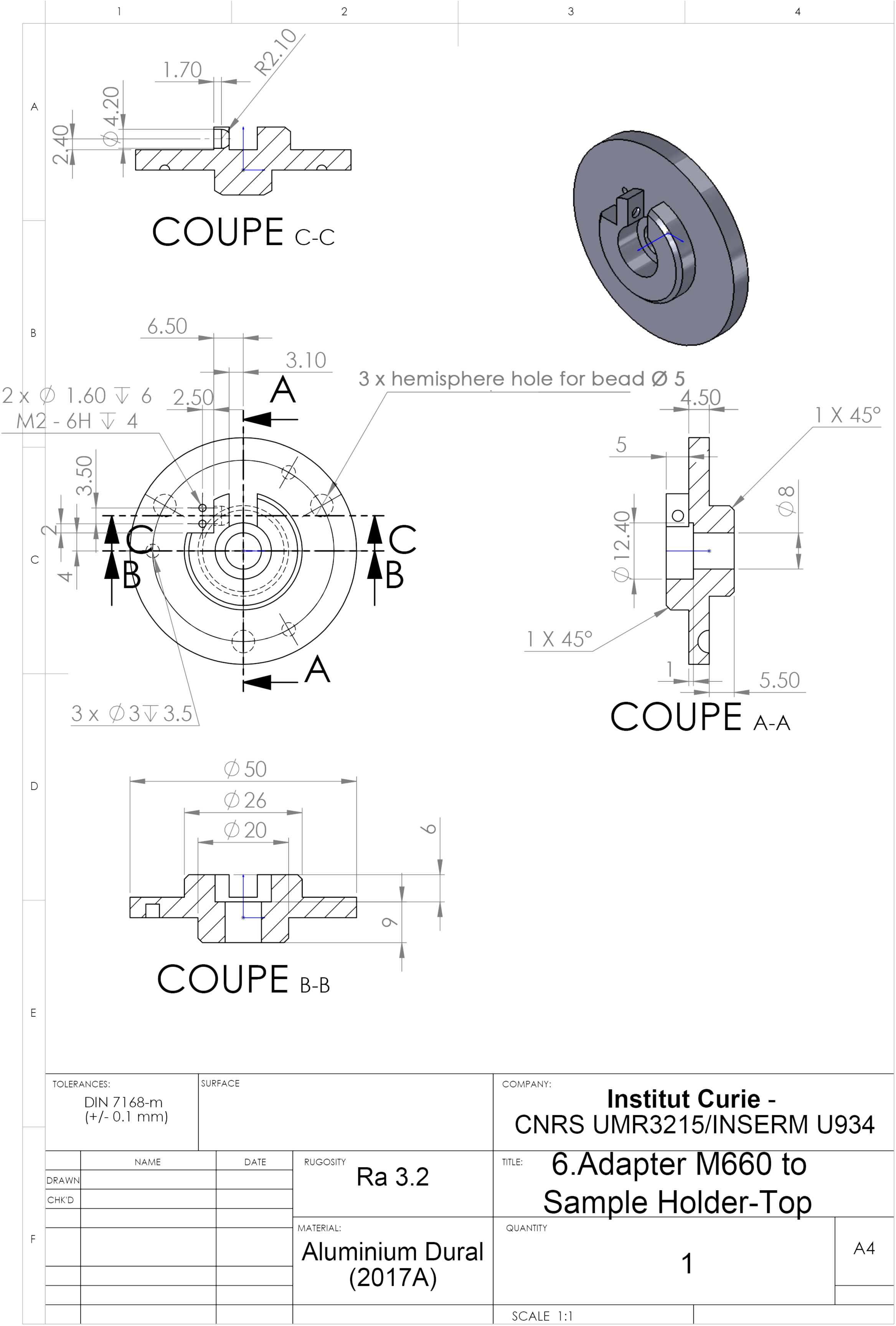
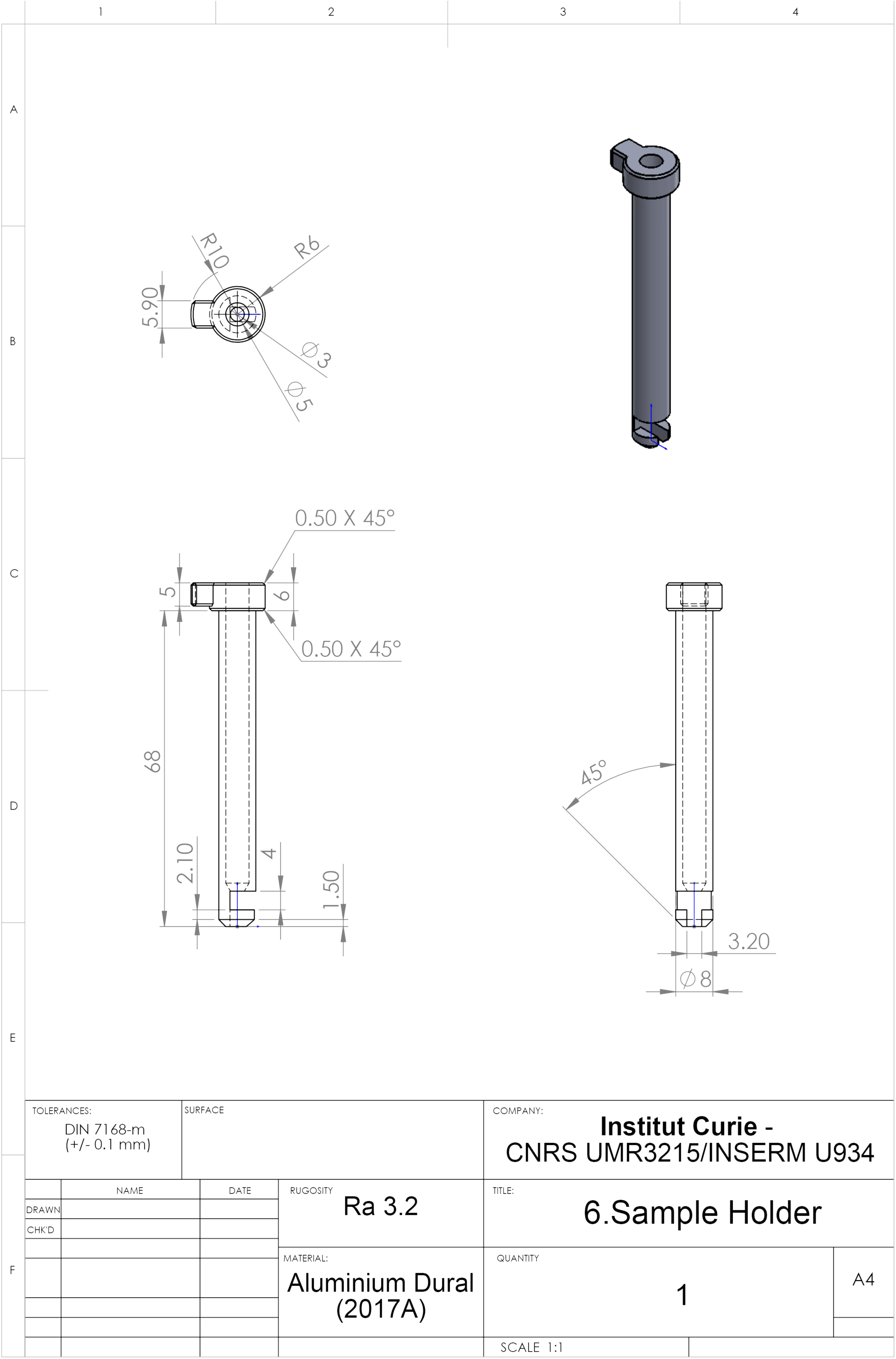
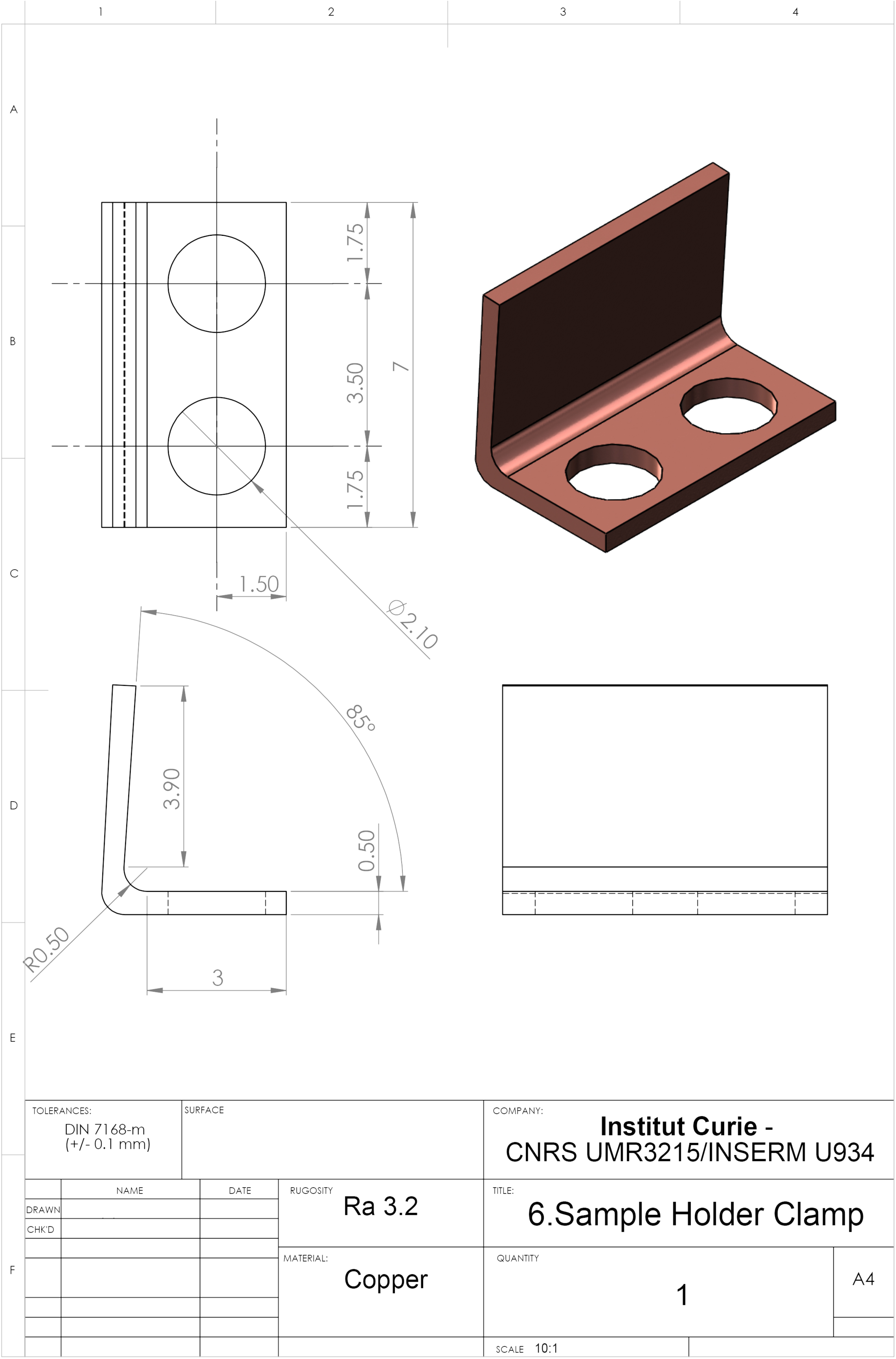
Blueprint of the Multi-View confocal microScope (MuViScope). A) Schematic illustration of the MuViScope showing the complete instrument that has been divided in eight blocks: (1) Camera and Spinning head, (2) Relay lens system, (3) Motorized flip mirror, (4) Right arm, (5) Left arm, (6) Specimen-positioning system, (7) Temperature and humidifier, (8) Computer and software. B) List of the optical elements, the opto-mechanical and the mechanical components used to mount the MuViScope. C) Technical drawings of the custom-made components: (Block 3) Scanner body, mirror body and mirror of the flip mirror; (Blocks 4 and 5) Adapters from SYS65 to SM2 and from SYS65 to SM1ZP; (Block 6) Adapters from M404 to M111, from M111 to M660 and from M660 to Sample Holder, Sample Holder and Clamp.

